# RRE-Finder: A Genome-Mining Tool for Class-Independent RiPP Discovery

**DOI:** 10.1101/2020.03.14.992123

**Authors:** Alexander M. Kloosterman, Kyle E. Shelton, Gilles P. van Wezel, Marnix H. Medema, Douglas A. Mitchell

## Abstract

Nearly half of the classes of natural products known as ribosomally synthesized and post-translationally modified peptides (RiPPs) are reliant on a protein domain called the RiPP recognition element (RRE) for peptide maturation. The RRE binds specifically to a linear precursor peptide and directs the post-translational modification enzymes to their substrate. Given its prevalence across various types of RiPP biosynthetic gene clusters (BGCs), the RRE could theoretically be used as a bioinformatic handle to identify novel classes of RiPPs. In addition, due to the high affinity and specificity of most RRE:precursor peptide complexes, a thorough understanding of the RRE domain could be exploited for biotechnological applications. However, sequence divergence of the RRE domain across RiPP classes has precluded automated identification of RREs based solely on sequence similarity. Here, we introduce RRE-Finder, a novel tool for identifying RRE domains with high sensitivity. RRE-Finder can be used in “precision” mode to confidently identify RREs in a class-specific manner or in “exploratory” mode, which was designed to assist in the discovery of novel RiPP classes. RRE-Finder operating in precision mode on the UniProtKB protein database retrieved over 30,000 high-confidence RREs spanning all characterized RRE-dependent RiPP classes, as well as several yet-uncharacterized RiPP, putatively novel gene cluster architectures that will require future experimental work. Finally, RRE-Finder was used in precision mode to explore a possible evolutionary origin of the RRE domain. Altogether, RRE-Finder provides a powerful new method to probe RiPP biosynthetic diversity and delivers a rich dataset of RRE sequences that will provide a foundation for deeper biochemical studies into this intriguing and versatile protein domain.

## Introduction

As of late 2019, nearly one-quarter of a million prokaryotic genomes were publicly available in the National Center for Biotechnology Information (NCBI) genome databases^1^. This vast genomic resource has been accelerating the pace of natural product discovery, with a recent surge of interest pertaining to the ribosomally synthesized and post-translationally modified peptides (RiPPs)^2^. RiPP biosynthesis starts with the ribosomal synthesis of a linear precursor peptide. The genes for RiPP precursor peptides are typically very short, have hypervariable sequences, and encode a peptide comprised of two parts—an N-terminal “leader” region and a C-terminal “core” region. With a few notable exceptions, the precursor peptide is genetically encoded adjacent to one or more genes encoding proteins that bind with high specificity and affinity to the leader region of the precursor. This interaction facilitates subsequent post-translational modification of the core residues^3^. After modification is complete, the leader peptide is often proteolytically removed and the mature RiPP product is exported from the producing organism^3^ (Figure 1). The exact nature of post-translational modifications is used to categorize RiPPs into individual classes, of which nearly 40 have been reported^2^. For example, lanthionine linkages define the lanthipeptide class while oxazol(in)e and thiazol(in)e heterocycles define the linear azol(in)e-containing peptide (LAP) class^4,5^.

**Figure 1.**
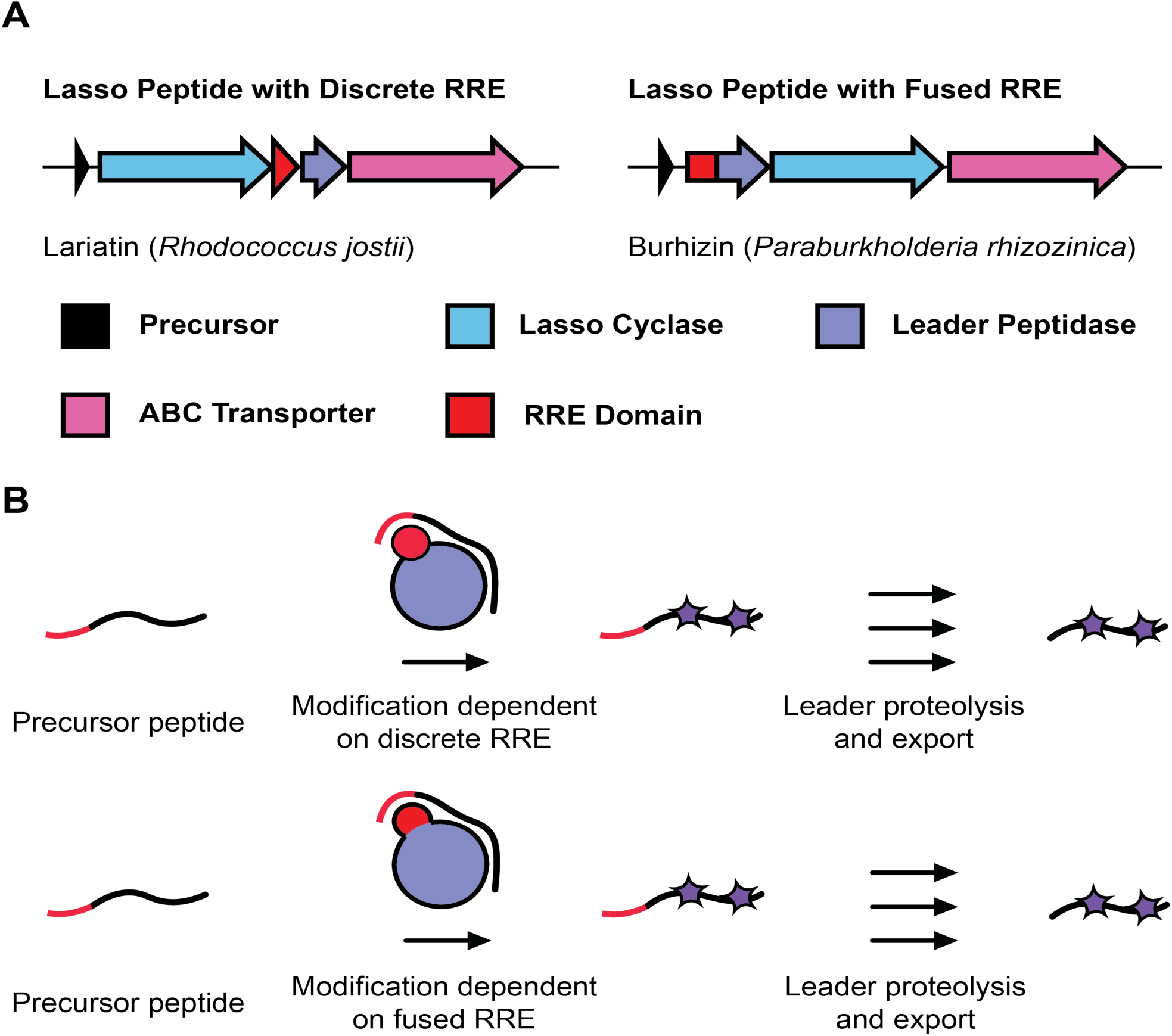
RRE-dependent RiPP biosynthesis. (A) RiPP biosynthetic gene clusters contain one or more short precursor peptides, often encoded adjacent to the modifying enzymes. Leader peptidases and proteins for immunity/export (often ABC transporters) are also frequently encoded in the local genome neighborhood. RRE domains are found fused to biosynthetic proteins as well as being produced as discrete proteins. (B) Modifying proteins bind the leader region of the precursor peptide employing the RRE domain. Post-translational modifications are then installed on the core region of the precursor peptide.

Many RiPP biosynthetic proteins recognize and bind their cognate precursor peptide through a domain known as the RiPP recognition element (RRE)^6^. The RRE consists of a conserved secondary structure of three, N-terminal alpha helices followed by a three-stranded beta sheet. The precursor peptide binds in a cleft between the third alpha helix (α3) and the third beta strand (β3), forming an ordered, four-stranded, antiparallel beta sheet (Figure S1). RRE domains can exist either as a discretely encoded protein (<100 residues) or as a fusion to a larger protein domain^6–10^. All RREs share sequence similarity to PqqD, which is a protein involved in pyrroloquinoline quinone (PQQ) cofactor synthesis—a redox cofactor produced by many prokaryotes^11^. Thus, the existence of a PqqD-like protein encoded nearby regulators, enzymes, and transporters is strongly indicative of an RRE-dependent RiPP cluster. The prevalence of PqqD-like proteins in RiPP clusters led to the discovery of the RRE domain and its conservation across RiPP classes in 2015^6^. Before this time, the importance of leader peptide recognition was established for a few RiPPs, such as nisin (lanthipeptide) and streptolysin S (LAP) biosynthesis^12,13^. In addition, an RRE-containing protein from microcin C7 synthesis (MccB) was co-crystallized with its cognate leader peptide in 2009, but it was not appreciated at the time that other RiPP classes employ a similar domain, due to RRE sequence divergence^14^.

Consistent with the rapid expansion of characterized RiPP BGCs, a diverse collection of modifications and enzymatic domains are found amongst the ∼40 known RiPP classes. However, the lack of a common genetic feature remains a major obstacle in the bioinformatic detection of novel RiPP classes. The fact that RRE domains are found in a major proportion of prokaryotic RiPP BGCs therefore provides a new opportunity: of the ∼30 known RiPP classes produced by prokaryotes, over 50% contain an identifiable RRE domain, which are found both as discrete polypeptides and as fusions to other biosynthetic proteins (Tables S1, S2). Considering the RRE domain appears to be the most conserved genetic feature found across prokaryotic RiPP classes, it theoretically could be used as an imperfect but useful bioinformatic handle to expand known RiPP sequence-function space by uncovering novel RRE-dependent RiPP classes.

The strategy outlined above is complicated by the sequence diversity of the RRE domain^6,9–11^. For example, if a pair-wise sequence alignment method (e.g. NCBI BLAST^16^) is used to compare RRE domains from two unrelated RiPP classes, sequence similarity will frequently not be detected, particularly in cases where the RRE domain is fused to a larger protein. The most appropriate Pfam^17^ model (a family of proteins sharing sequence homology) for defining the RRE domain is PF05402, which extensively covers bona fide PqqD proteins from PQQ BGCs. PF05402 incompletely retrieves RRE-containing proteins from only a few other RiPP classes (e.g. lasso peptides and sactipeptides) and most RRE-dependent RiPP classes are not represented in this Pfam^18,19,20^ (Figure S2). These results underscore the inability of a single bioinformatic model to capture the breadth of RRE sequence diversity. Owing to the fact that RREs share considerable structural similarity, HHpred^21^ is a more sensitive algorithm for detecting RRE domains. HHpred detects remote protein homology by aligning profile hidden Markov models (pHMMs; a model that defines amino acid frequency for a protein family) and comparing their (predicted) secondary structures. RREs were originally detected using this method by analyzing several RiPP modifying enzymes, which showed consistent homology to PqqD^6^. However, HHpred requires generation of a multiple sequence alignment (MSA) and secondary structure prediction using PSIPRED^22^. These steps require several minutes of computing time per protein query, rendering the process unattractive for large datasets. In this work, we report a customized tool that permits the rapid and accurate detection of RREs in known and potentially novel RiPP classes with the principle goal of directing natural product hunters to the most fruitful areas of the RiPP sequence-function space.

## Results and Discussion

### Development of RRE-Finder

This work presents a new tool for mining microbial genomes in search of RRE domains, called RRE-Finder. This tool has two modes of operation. The first is “precision” mode, which employs a set of 35 custom pHMMs designed to detect RRE domains in a class-dependent manner (Figure S3, Table S3 Dataset S1). The precision mode pHMMs are primarily based on known RiPP classes—in most cases, representative RRE-containing proteins from these classes have been verified to bind their cognate precursor peptide either through X-ray crystallography or binding assays such as fluorescence polarization. The second mode, “exploratory” mode, uses a truncated version of the HHpred^21^ pipeline with a custom database of detected RREs. Depending on the end-user’s objective, RRE-Finder can be used in precision mode to accurately predict the presence of an RRE domain as well as the likely RiPP class in which the precursor peptide belongs. Alternatively, in exploratory mode, the user can retrieve a wider array of putative RRE-containing proteins to assist in the discovery of novel RRE-dependent RiPP classes. RRE-Finder accelerates the process of identifying RRE domains by several orders of magnitude compared to HHPred. Precision mode, for instance, can analyze >5,000 protein sequences per second (Table S4).

In addition to models based on known RiPP classes, precision mode includes several “auxiliary” models based on high confidence, novel RiPP classes. We justified the inclusion of these models based on repeated observation of RRE domains within RiPP-like genomic contexts across multiple prokaryotic species. In general, for RiPP classes where an extensive survey of the bioinformatic space has been performed (e.g. lasso peptides^23,24^, sactipeptides and rantipeptides^25^, thiopeptides^26^), custom HMMs were built by first visualizing sequence space through use of a sequence similarity network (SSN) for all RRE-containing proteins in the dataset^27^. SSN visualization using the Cytoscape tool^28^ facilitated selection of the most diverse and nonredundant subset of RRE primary sequences for MSA seed sequence alignment. In cases where a published dataset was available for a given RiPP class, model prediction accuracy was gauged by using HMMscan (from the HMMER3 suite of tools^29^) on the relevant dataset using bit score cutoffs of 15, 25, and 35 (subsequently referred to as tolerant, moderate, and stringent cutoffs). A given HMM was considered acceptable if >95% of RRE-containing proteins within the dataset were retrieved by the model at a bit score of 25 (Table S5).

In cases where a deep, bioinformatic profiling of a RiPP class has not been previously published or where a mature natural product is not known (i.e. for generating the auxiliary models), seed alignment input sequences were gathered using PSI-BLAST^30^ to find diverse homologous sequences to a representative sequence from each given class. The generated HMMs were considered valid if an HMMsearch of the UniProtKB database^31^ with a bit score cutoff of 25 gave only hits with similar BGC architecture to the target class. In addition, characterized datasets of RiPP proteins (e.g. lanthipeptides^32,33^, lasso peptides^23,24^, and sactipeptides^25^) were used to test auxiliary models using HMMscan analysis. Models giving few or no hits were considered to have acceptably low false positive rates.

Exploratory mode, on the other hand, was built for the detection of RRE domains with greater sequence divergence from those detected by precision mode. For this mode, we employed a variation of the HHpred pipeline to detect structural similarity to RRE domains. The HHpred pipeline uses a clustered UniProt database (uniclust30)^34^, which comprises a small, representative set of all UniProt protein sequence diversity. Query proteins are compared to the uniclust30 database to generate a representative protein family for the query, and the consensus sequence of this representative protein family is compared to those of other protein families. This search also incorporates comparison of (predicted) secondary structures. As such, HHpred can detect distantly related sequences and overlap in secondary structures between a query protein and the UniProt database. However, the vast search space used far exceeds what is necessary if the only goal is to detect RRE domains.

To accelerate the HHpred pipeline for RRE detection, we first built a smaller, more specialized HHpred database, consisting of ∼3,400 diverse RRE sequences. These sequences were gathered by mining 5,000 RiPP BGCs from the antiSMASH database^35^ using HHpred. Rather than manually curating the retrieved RREs in a class-specific manner, as was done for precision mode, we included all detected RREs indiscriminately. The selected RREs were supplemented with 7 RREs from LAP BGCs and one RRE from a proteusin BGC, as no BGCs from these RiPP classes were present in the antiSMASH database.

The collection of 3,413 RREs was used to build databases for two filtering steps (Figure 2). For the first filter, all RREs were clustered into representative protein families with MMSeqs2^36^, resulting in 558 RRE families. These RRE families were further enriched by querying each family against the uniclust30 database using hhblits, an iterative search tool from HHpred^37^. For each of the 558 RRE families, custom pHMMs were constructed, allowing for an initial filtering step with HMMsearch^29^. The second filtering step functions in a similar manner to HHpred. However, rather than using the uniclust30 database to retrieve a protein family for a query, we employed a smaller, custom HHpred database consisting of the 3,413 RRE sequences retrieved from the antiSMASH database and additional sequences retrieved by hhblits. Using this custom database, only protein queries that are homologous to one of the 558 clustered RRE families will return results. For queries without homology, no protein family would be found in the database, effectively filtering out these sequences. Finally, exploratory mode compares the family of proteins homologous to a query protein to three RRE structures in the Protein Data Bank (PDB entries: 5V1T, 5SXY, 3G2B). Any proteins showing homology to these models are output as putative RRE domains. In all, by employing a small, custom library of RRE sequences, exploratory mode significantly accelerates detection of RREs over the HHpred pipeline.

**Figure 2.**
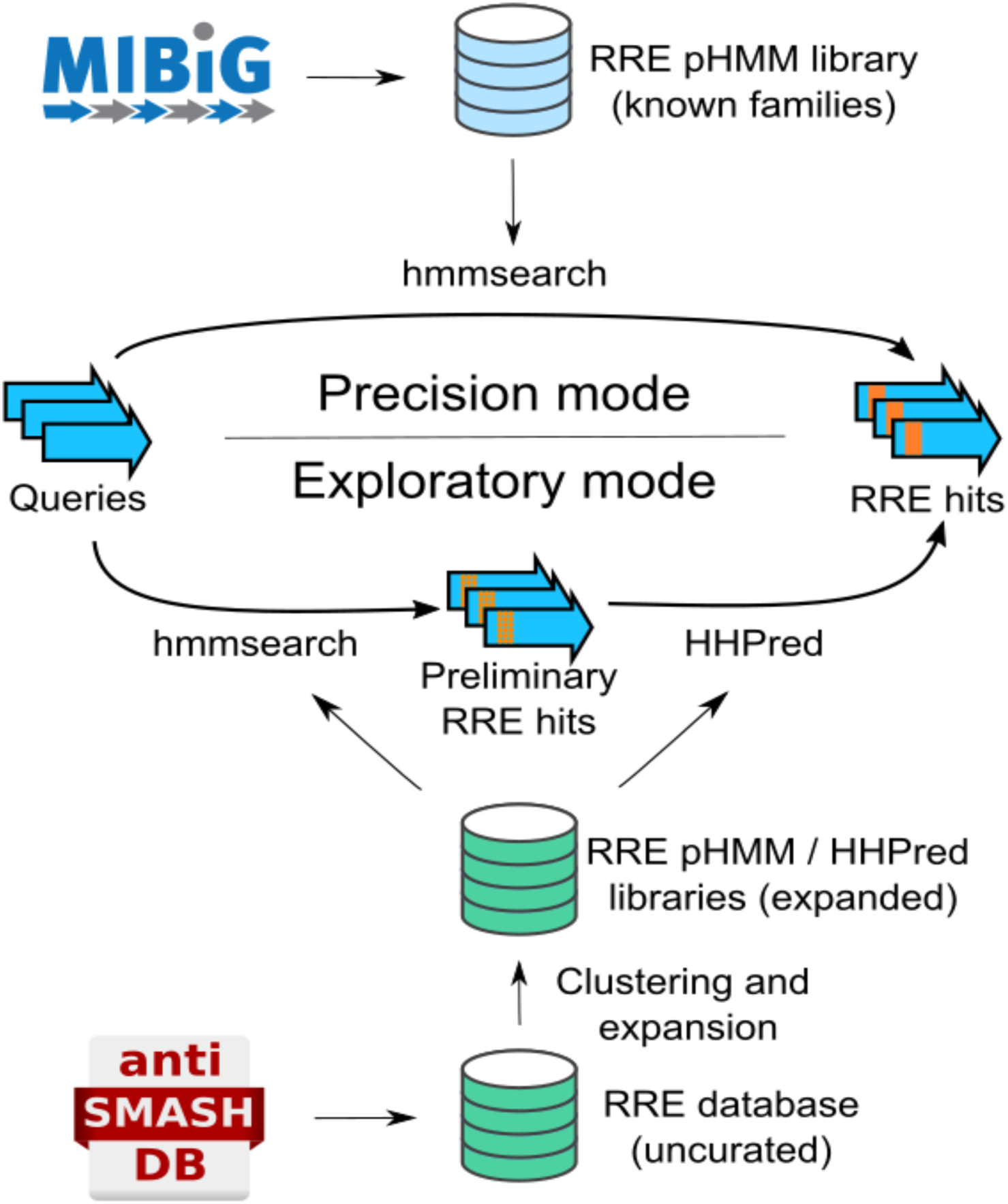
RRE-Finder employs two modes for RRE detection. Precision mode (top) of RRE-Finder uses a set of pHMMs to accurately predict RREs. These pHMMs are based on characterized RRE domains for individual RiPP classes, either from published datasets or from the MIBiG database. Exploratory mode uses a combination of pHMMs and a truncated HHPred pipeline (including secondary structure prediction) to identify divergent RRE sequences (albeit with a higher false-positive rate).

### Model Validation Against the MIBiG Database

As an initial test of accuracy, RRE-Finder was evaluated in precision and exploratory modes against the MIBiG database^38^. This database contains characterized BGCs for almost 2,000 natural products, including polyketides, non-ribosomal peptides, and RiPPs. All proteins within the MIBiG set (version 1.4) of RiPP (n = 242) and non-RiPP BGCs (n = 1,575) were analyzed by RRE-Finder at tolerant, moderate, and stringent bit score cutoffs (Figure 3).

**Figure 3.**
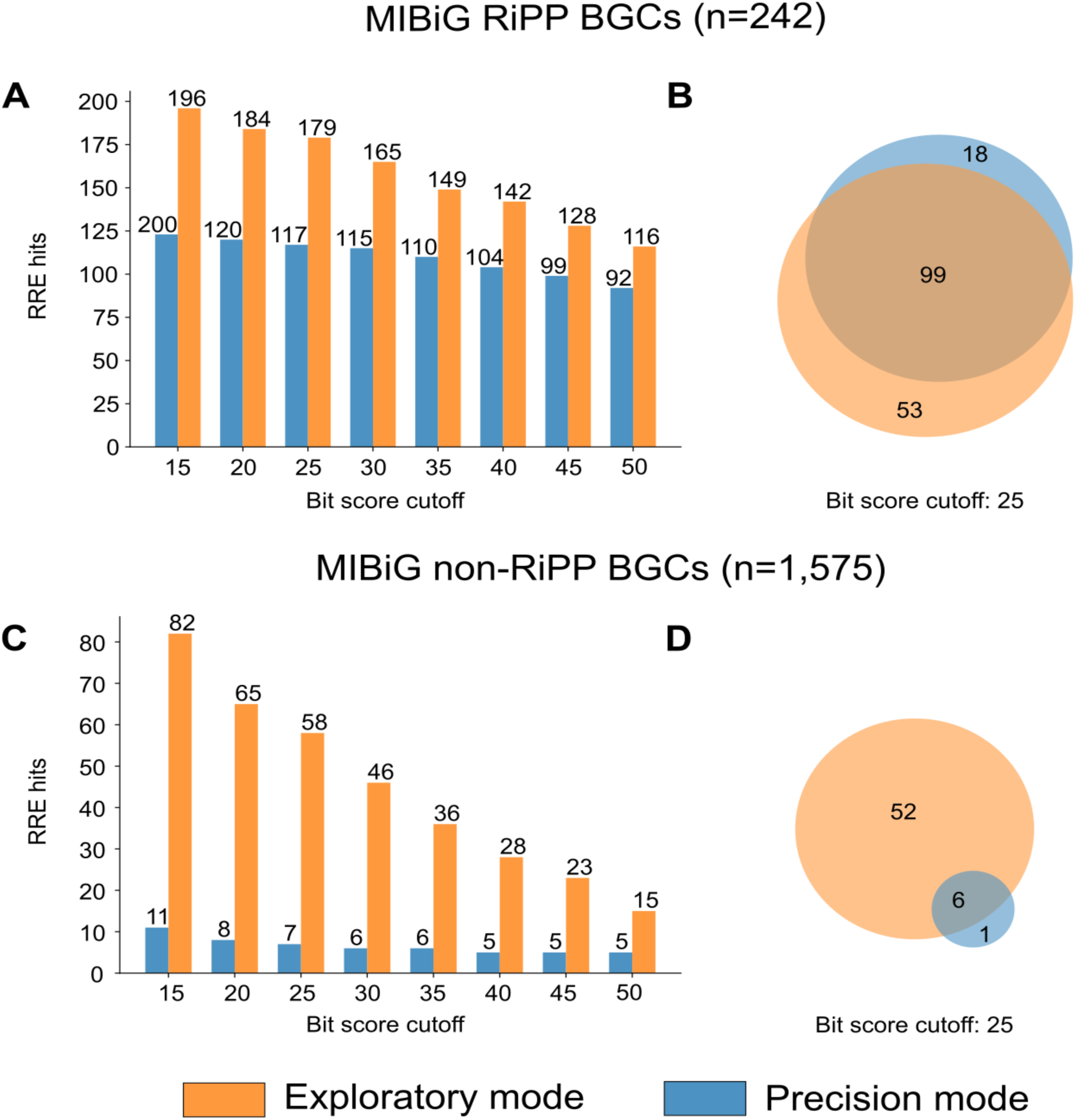
MIBiG validation of RRE-Finder. Both modes were used to retrieve RRE-containing proteins in 242 RiPP BGCs (A, B) and 1,575 non-RiPP BGCs (C, D) from the MIBiG database. With increasing bit score stringency, the number of RRE detected decreased in both types of BGCs (A, C). At a bit score of 25, exploratory mode of RRE-Finder detects most of the RREs found by precision mode in RiPP BGCs (B), as well as several other RREs. However, the number of RREs detected in non-RiPP BGCs is significantly lower for precision mode compared to exploratory mode (D).

In general, both precision and exploratory modes accurately predicted the presence of RRE domains in >90% of the RRE-dependent RiPP BGCs. Taken together, both modes retrieved 93% (115/122) of known RRE-containing proteins (Table S6). With increasing bit score stringency, the number of RRE sequences retrieved decreased in both RiPP and non-RiPP BGCs, as expected (Figure 3). At all bit score cutoffs, exploratory mode predicted more RRE domains in RiPP BGCs (higher true positive rate compared to precision mode), while precision mode retrieved fewer proteins from non-RiPP BGCs (lower false positive rate compared to exploratory mode).

Further analysis of the results at varied bit score stringencies led us to choose a bit score cutoff of 25 as a middle ground between precision and recall. At this cutoff, most of the RREs found within the MIBiG set by precision mode were also found by exploratory mode (101/117, Figure 3). Only the RREs of linear-azol(in)e containing peptides (LAPs)^4^, streptides^39^, and polytheonamides^40^ proved more difficult to detect by exploratory mode (Table S6). Further enriching the exploratory mode training set with RREs from these RiPP classes restored detection of the RREs from the polytheonamide cluster, although precision mode proved most consistent for RRE detection within the LAP and streptide classes.

In contrast, precision mode detected only 66% (101/152) of the RREs retrieved by exploratory mode. A significant number (n = 18) of the RRE-containing proteins not detected by precision mode were those contained in LanB-like proteins, which are found in lanthipeptide and thiopeptide BGCs. It has been shown that the LanB RRE domain found in thiopeptide BGCs is possibly vestigial as the cognate leader peptide is not required for catalytic processing^41^. Exploratory mode also detected a significant number (n = 14) of RREs fused to dehydrogenase enzymes present in cyanobactin, LAP, and thiopeptide BGCs, which were not detected by precision mode. These RREs are also perhaps vestigial; thus, precision mode does not include a model for identifying this type of RRE domain.

While exploratory mode detects a greater number of RREs, it also displays a higher false positive rate (e.g. proteins retrieved from non-RiPP clusters). These retrieved sequences primarily consisted of helix-turn-helix domains, methyltransferases, and proteins with homology to known RRE-containing proteins that occur in non-RiPP contexts, such as radical *S*-adenosylmethionine (rSAM) enzymes (Table S7). Many DNA-binding regulators possess a helix-turn-helix domain, which are structurally homologous to RRE domains (Figure S4). Indeed, most RRE domains analyzed by HHpred show homology to known DNA-binding domains and regulatory elements (e.g. PDB entries: 3DEE, 2G9W, 2OBP). Because regulatory proteins are not known to bind or modify RiPP precursor peptides, RRE-Finder includes an option to filter out proteins that correspond to such domains.

RRE-Finder operating in either mode retrieved LanB-like proteins within polyketide BGCs. There is precedence for the assimilation of RiPP-modifying enzymes into polyketide pathways^32^, although the RRE domain within these proteins may be vestigial (Figure S5). Thus, retrieval of proteins outside of canonical RiPP BGCs may not be false positives. Further biochemical validation is required to confirm or refute a functional RRE in these instances.

Finally, some HMMs employed by precision mode were generated largely using RRE sequences from the MIBiG database. In these cases, validation against MIBiG alone is not sufficient to prove these models exhibit appropriate recall and precision. As an orthogonal means of precision mode validation, we performed an HMMscan of the 5,000 RiPP BGCs from the antiSMASH database initially used to generate the exploratory mode custom databases^35^. As previously stated, these BGCs primarily belong to the lanthipeptide, thiopeptide, LAP, sactipeptide, and lasso peptide classes. Because this collection of gene clusters was not curated to include only RRE-dependent RiPP BGCs, there are clusters not anticipated to be retrieved by precision mode (e.g. class II-IV lanthipeptides)^15^. These clusters were purposely included in the analysis as a negative control. All proteins within each of the 5,000 BGCs were scanned by precision mode at tolerant, moderate, and stringent bit score cutoffs. At these bit score stringencies, the percentage of gene clusters predicted by precision mode to contain an RRE were 90%, 87%, and 83%, respectively. The 10% of BGCs not predicted to contain an RRE by precision mode were manually examined, and the majority of these clusters belong to RiPP classes that are RRE-independent. Thus, precision mode accurately predicts the presence of RREs in an unbiased collection of gene clusters, and appropriately filters out RRE-independent RiPP clusters.

### Defining the Scope of RRE-Dependent RiPP BGCs

Next, we aimed to profile the extent to which the RRE domain is present within sequenced genomes by mining the entire UniProtKB database with both modes of RRE-Finder^31^. Using HMMsearch at a moderate bit score cutoff of 25, precision mode retrieved ∼30,000 proteins (∼13,000 non-redundant sequences, Figure 4). A parallel search using exploratory mode yielded ∼70,000 non-redundant RRE-containing proteins, almost completely encompassing the proteins retrieved by precision mode, except for three proteins. As expected, the numbers of proteins retrieved by precision mode is larger than has been previously reported for virtually all RiPP classes, owing to on-going genome sequencing. For example, the thiopeptide F protein model is the top-scoring model for ∼500 of the retrieved UniProtKB proteins, which roughly a 25% increase from the most recent bioinformatic survey of thiopeptide BGCs^26^. In some cases, however, the number of retrieved proteins for a given model may be misleading. For example, the precision mode model specific to discretely encoded lasso peptide RREs is the top-scoring model for ∼5,000 of the retrieved proteins. However, co-occurrence analysis of these clusters revealed that only ∼3,500 of the retrieved proteins co-occur with the expected leader peptidase and lasso cyclase enzymes. This number is more consistent with the most recent bioinformatic survey of lasso peptides, which reported ∼3,000 lasso peptide BGCs with discretely encoded RREs^24,42^. Proteins retrieved by this model often co-occur with other common RiPP enzymes, such as rSAM enzymes (which represent ∼300 of the “false positive” BGCs). Thus, we caution that the number of proteins retrieved by any given model should not be equated to the number of gene clusters specific to a particular RiPP class without analysis of the local genomic neighborhood.

**Figure 4.**
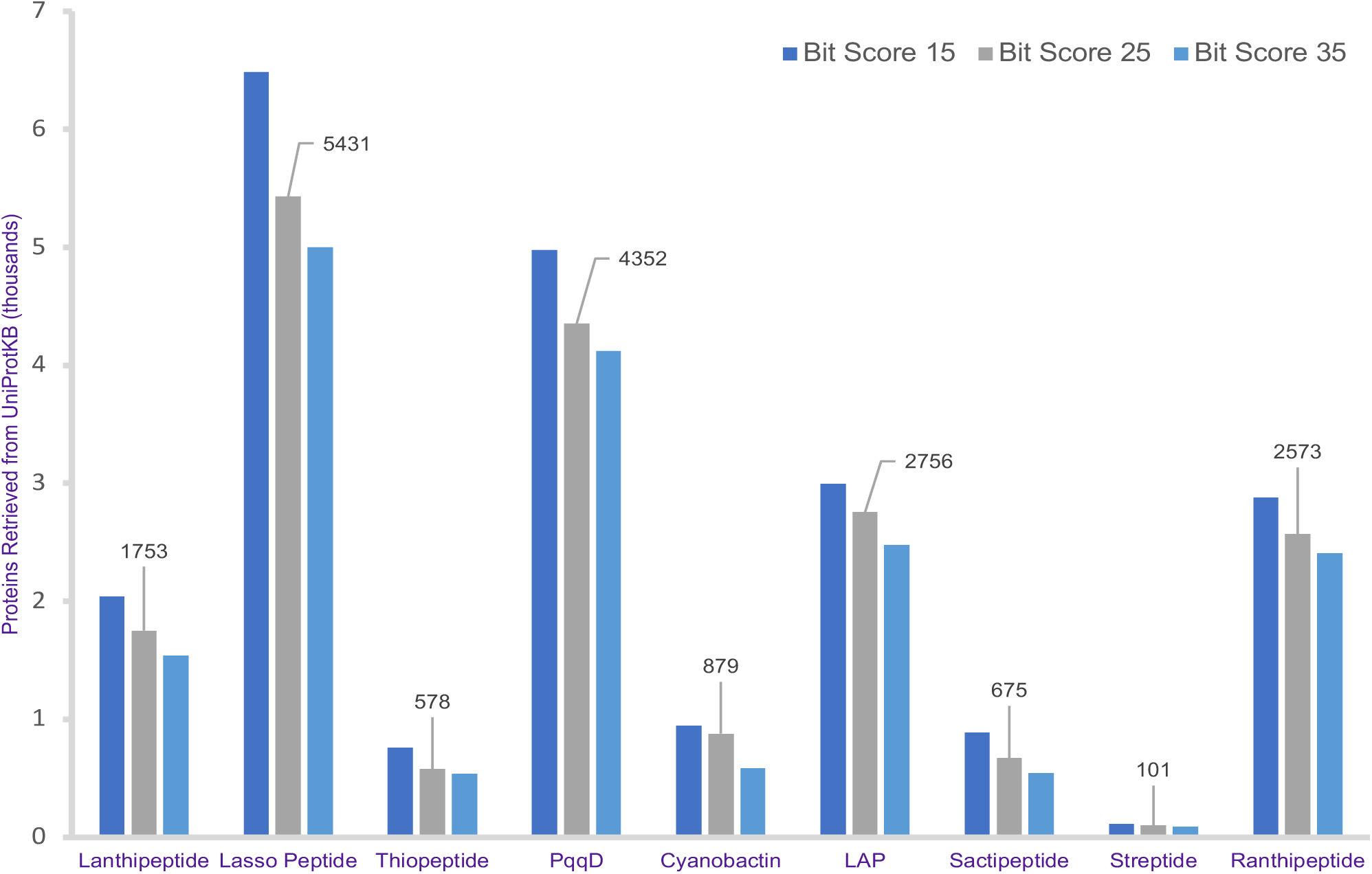
Summary of precision mode model hits to UniProtKB. The number of protein matches within the UniProt database are summarized for several large classes of RiPPs. A scan of the entire UniProt database of non-redundant proteins was carried out at three separate bit score cutoffs. Number of hits for a bit score 25 cutoff is indicated above the data for each class. Full data on hits to each precision mode model are available in Supplementary Dataset 1.

The excised RRE sequences from all proteins identified by precision mode were visualized using a sequence similarity network (SSN)^27^. This SSN confirms known relationships between RREs in separate RiPP classes. For example, discretely encoded lasso peptide RREs (referred to as either the B1 or E protein) group separately from RRE-peptidase fusions (known as the B protein), consistent with a different recognition sequence on the leader peptide for these two varieties of lasso peptides (Figures 5, S6, S7)^23,24^. In contrast, the heterocycloanthracins (a subclass of LAPs), cluster more tightly with thiopeptides than they do other LAPs. This relationship was expected given that heterocycloanthracin and thiopeptide BGCs feature an RRE domain fused to an “ocin-ThiF-like” protein (TIGR0393) that delivers the peptide substrate to the biosynthetic enzymes^4,43^. Other LAP pathways do not fuse the RRE domain to a member of TIGR0393 but rather contain an RRE fusion to members of TIGR03882^4,6,43,44^. Another method to view RRE relatedness is through model redundancy (Figures S8, S9). In cases where there is significant overlap in the proteins retrieved by one or more models (i.e. thiopeptides and heterocycloanthracins; goadsporins and cyanobactins), this model redundancy is reflective of RREs in these classes binding their cognate leader peptides through similar motifs. Similarly, lack of model overlap is indicative of a divergent leader peptide recognition sequence. For example, even at a tolerant bit score cutoff of 15, there is virtually no overlap between the lanthipeptide-associated RRE domains with any other RiPP class, reflective of a unique recognition sequence not observed in other classes^45,15^ (Figure S8).

**Figure 5.**
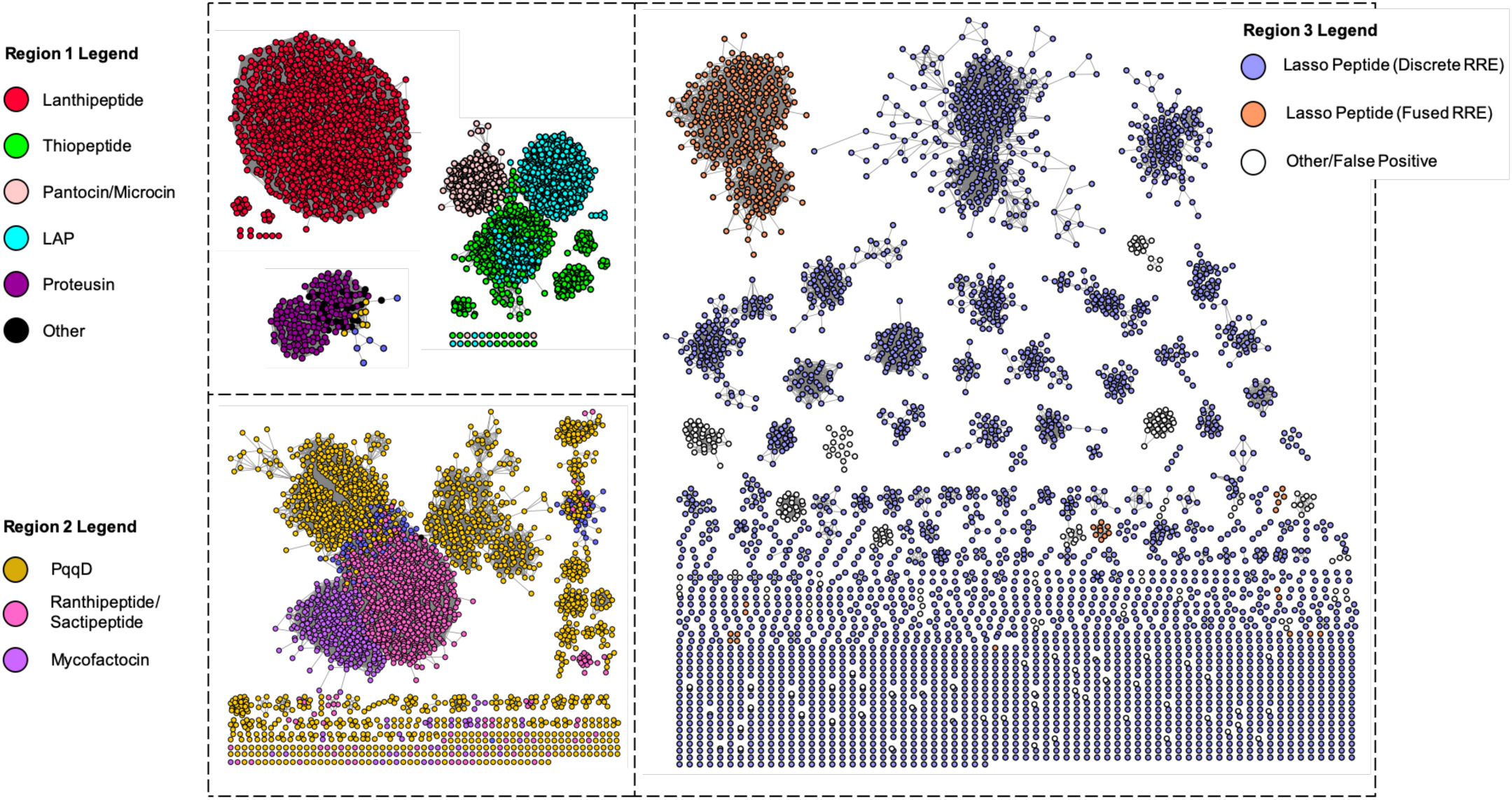
Sequence similarity network of UniProtKB proteins retrieved by precision mode. Shown is a RepNode60 SSN at an alignment score of 22 (sequences with >60% identity are conflated to a single node and edges represent a BLAST expectation value better than 10^−22^). Proteins are colored based on the best-fit model by which they were detected. White nodes in region 3 represent proteins that were retrieved by the discrete lasso peptide RRE model but do not co-occur with the requisite lasso peptide machinery (i.e. a leader peptidase and a lasso cyclase). These proteins represent possible false positives from this model. The discrete lasso peptide RREs clustering with sacti/ranthipeptides in region 2 are stand-alone RRE proteins that co-occur with radical SAM enzymes. The network was generated using the EFI-EST webtool (https://efi.igb.illinois.edu/efi-est/)^27^.

### Evolution of the RRE Domain

Sequence similarity between leader peptide recognition sequences in closely related RiPP classes suggests that the RRE domain emerged once and then diverged to recognize a variety of leader peptides. Because the leader peptide binds as an ordered beta-strand between the α3 helix and β3 strand of the RRE, substitution of key α3 and β3 residues logically tune the RRE specificity to target peptides. In fact, analysis of residue-level conservation between RREs of divergent RiPP classes reveals that the α3 and β3 regions exhibit higher levels of residue conservation than the rest of the RRE, presumably due to selective pressure to conserve leader peptide:RRE contacts. This remains true when comparing closely related RiPP classes, such as LAPs and thiopeptides (Table S8). The other regions of the RRE, which are not directly involved in binding the leader peptide, show much lower levels of conservation when compared to the α3 and β3 regions; these regions are more tolerant of mutation, so long as the α3-β3 secondary structure is maintained.

A representative diversity-maximized phylogenetic tree of excised RRE domains retrieved by precision mode (bit score cutoff of 25) supports the hypothesis that the RRE domain co-evolved with the leader peptide to expand specificity to all RRE-dependent RiPP classes (Figure S10). The sequences contained within this tree do not include all proteins retrieved by precision mode from the UniProtKB database. Instead, 10% of the proteins contained within each cluster of the SSN (Figure 5) were employed, along with all singleton nodes, to generate a representative collection of sequences spanning all RRE-dependent classes. The tree employs a helix-turn-helix DNA-binding protein as an outgroup (PDB entry 3DEE), as this protein scores well in HHpred searches of characterized RRE proteins, such as PqqD and LynD. As previously mentioned, it is plausible that the RRE domain evolved from DNA-binding regulatory elements, given the shared secondary structure and the similar function of these domains to specifically bind a stretch of DNA or a peptide (Figure S4). Unsurprisingly, the diversity-maximized tree shows a closer relationship between the helix-turn-helix outgroup and discretely encoded RRE domains, such as in PQQ and lasso peptide clusters. This is consistent with the theory that stand-alone RRE domains were likely the first types of RRE domains to evolve. Furthermore, all fused RREs contained within the tree form monophyletic clades, suggesting that fused RREs evolved separately for separate RiPP classes. Once again, this is unsurprising as some classes employ N-terminal fused RRE domains, while others code for C-terminal (e.g. proteusins) or internal RREs (e.g. lanthipeptides).

### Using RRE-Finder to Identify Novel RiPP Clusters

Theoretically, the sequence space retrieved by exploratory mode and the auxiliary models of precision mode encompasses RRE-containing proteins from potentially novel RiPP classes. To explore this sequence space, divergent clusters mined from UniProtKB were manually examined for novel RiPP contexts. All proteins retrieved were grouped based on their best fit Pfam model. As expected, many of the proteins retrieved by exploratory mode are clear false positives—either regulatory elements or other proteins containing helix-turn-helix motifs. The proteins retrieved also included 1,050 methyltransferases, which were assumed to be false positives, given their prevalent occurrence in non-RiPP BGCs (Table S9).

Excluding these false positives, RRE-Finder reveals many potentially novel RiPP clusters with new gene architectures containing both discrete and fused RRE domains. Included in these clusters are RRE-protein fusions that are not present in known classes, such as RRE-glycosyltransferase fusions and RRE-glutathione *S*-transferase fusions (Figure S11, Table S10). Of the nine potential RiPP clusters shown in Figure S11, four contain genes encoding rSAM enzymes, which are common in several RiPP classes^25^. The presence of rSAM enzymes in conjunction with predicted RREs is suggestive of a RiPP BGC. However, out of the nine gene clusters, only three contained probable precursor peptides (small genes <150 amino acids, clustered with the RRE-containing protein); therefore, manual curating of potentially novel clusters found by RRE-Finder is recommended.

### RRE-Finder incorporation into antiSMASH and RODEO

To democratize the use of RRE-Finder to identify RREs in gene clusters of interest, RRE-Finder is available as a standalone command-line tool (https://github.com/Alexamk/RREFinder). Protein queries can be input in either FASTA or GenBank formats, and the tool is also capable of analyzing and updating antiSMASH output files. Precision mode of RRE-Finder will be incorporated into the next release of antiSMASH. We have also incorporated the precision models of RRE-Finder into RODEO^23^, which is a genome-mining tool for RiPP discovery that provides genomic neighborhood visualization and prediction of precursor peptides. Protein-coding sequences within the genetic locus are annotated according to Pfam and TIGRFAM models to identify conserved domains and predict function. With the “include RRE scoring” function enabled, proteins that contain an identifiable RRE are annotated, along with their E-value significance. In cases of fused RRE domains, the position of the RRE domain within the larger protein is specified. RRE predictive scoring, powered by RRE-Finder, is now available within the command line (https://github.com/the-mitchell-lab/rodeo2) and webtool (http://rodeo.scs.illinois.edu) versions of RODEO.

## Conclusion

RRE-Finder rapidly and accurately detects RRE domains within known and potentially novel RiPP classes. Although not all RiPP classes are RRE-dependent, the majority of prokaryotic RiPP classes are, including the largest known classes (i.e. class I lanthipeptides, lasso peptides, and ranthipeptides). RiPP natural products are a prime candidate for pathway engineering, as precursor peptides and their cognate modifying enzymes are all genetically encoded, typically within one BGC. However, efforts to bioinformatically predict RiPP BGCs lag behind those for predicting PKS/NRPS clusters, due to a lack of strongly conserved protein domains spanning multiple RiPP classes. Virtually no novel RiPP classes have been discovered using solely a bioinformatic approach: The ranthipeptide class was defined solely using bioinformatics, however this class had previously been predicted and misclassified as sactipeptides^25^. Furthermore, bioinformatics have been used to vastly expand known diversity within some classes—for example, the streptide class has been expanded to include enzymes that diverge from the class-defining Lys-Trp crosslinking enzymes^39,46^. Recently, two new RiPP classes—the peptide amino-acyl tRNA ligases (PEARLs) and the α-keto β-amino acid-containing peptides—were discovered through bioinformatic means^47,48^. However, both of these classes were discovered through the presence of known RiPP biosynthetic enzymes within the clusters, rather than through unbiased bioinformatic discovery. Through precision mode of RRE-Finder, we have shown that characterized RiPP classes contain more members than currently reported, although analysis of the genomic neighborhood is often necessary to confirm class identity. Precision mode can further be employed, particularly with low bit score stringency, to predict novel RRE domains, such as those predicted by the auxiliary models. Finally, using RRE-Finder in exploratory mode reveals a set of over 70,000 proteins that are predicted to contain an RRE, suggesting that many more novel classes of RRE-dependent RiPPs are likely to exist in nature.

## References

(1) Ncbi Resource Coordinators. Database Resources of the National Center for Biotechnology Information. Nucleic Acids Res 2015, 43, D6–17. https://doi.org/10.1093/nar/gku1130.

(2) Arnison, P. G.; Bibb, M. J.; Bierbaum, G.; Bowers, A. A.; Bugni, T. S.; Bulaj, G.; Camarero, J. A.; Campopiano, D. J.; Challis, G. L.; Clardy, J.; Cotter, P. D.; Craik, D. J.; Dawson, M.; Dittmann, E.; Donadio, S.; Dorrestein, P. C.; Entian, K. D.; Fischbach, M. A.; Garavelli, J. S.; Goransson, U.; Gruber, C. W.; Haft, D. H.; Hemscheidt, T. K.; Hertweck, C.; Hill, C.; Horswill, A. R.; Jaspars, M.; Kelly, W. L.; Klinman, J. P.; Kuipers, O. P.; Link, A. J.; Liu, W.; Marahiel, M. A.; Mitchell, D. A.; Moll, G. N.; Moore, B. S.; Muller, R.; Nair, S. K.; Nes, I. F.; Norris, G. E.; Olivera, B. M.; Onaka, H.; Patchett, M. L.; Piel, J.; Reaney, M. J.; Rebuffat, S.; Ross, R. P.; Sahl, H. G.; Schmidt, E W.; Selsted, M. E.; Severinov, K.; Shen, B.; Sivonen, K.; Smith, L.; Stein, T.; Sussmuth, R. D.; Tagg, J. R.; Tang, G. L.; Truman, A. W.; Vederas, J. C.; Walsh, C. T.; Walton, J. D.; Wenzel, S. C.; Willey, J. M.; van der Donk, W. A. Ribosomally Synthesized and Post-Translationally Modified Peptide Natural Products: Overview and Recommendations for a Universal Nomenclature. Natural product reports 2013, 30, 108–160. https://doi.org/10.1039/c2np20085f.

(3) Hudson, G. A.; Mitchell, D. A. RiPP Antibiotics: Biosynthesis and Engineering Potential. Curr Opin Microbiol 2018, 45, 61–69. https://doi.org/10.1016/j.mib.2018.02.010.

(4) Cox, C. L.; Doroghazi, J. R.; Mitchell, D. A. The Genomic Landscape of Ribosomal Peptides Containing Thiazole and Oxazole Heterocycles. BMC genomics 2015, 16, 778. https://doi.org/10.1186/s12864-015-2008-0.

(5) Zhang, Q.; Yu, Y.; Velasquez, J. E.; van der Donk, W. A. Evolution of Lanthipeptide Synthetases. Proc Natl Acad Sci USA 2012, 109 (45), 18361–18366. https://doi.org/10.1073/pnas.1210393109.

(6) Burkhart, B. J.; Hudson, G. A.; Dunbar, K. L.; Mitchell, D. A. A Prevalent Peptide-Binding Domain Guides Ribosomal Natural Product Biosynthesis. Nat. Chem. Biol. 2015, 11, 564–570. https://doi.org/10.1038/nchembio.1856 http://www.nature.com/nchembio/journal/v11/n8/abs/nchembio.1856.html#supplementary-information.

(7) Davis, K. M.; Schramma, K. R.; Hansen, W. A.; Bacik, J. P.; Khare, S. D.; Seyedsayamdost, M. R.; Ando, N. Structures of the Peptide-Modifying Radical SAM Enzyme SuiB Elucidate the Basis of Substrate Recognition. Proceedings of the National Academy of Sciences of the United States of America 2017, 114, 10420–10425. https://doi.org/10.1073/pnas.1703663114.

(8) Ortega, M. A.; Hao, Y.; Zhang, Q.; Walker, M. C.; van der Donk, W. A.; Nair, S. K. Structure and Mechanism of the TRNA-Dependent Lantibiotic Dehydratase NisB. Nature 2014. https://doi.org/10.1038/nature13888.

(9) Koehnke, J.; Mann, G.; Bent, A. F.; Ludewig, H.; Shirran, S.; Botting, C.; Lebl, T.; Houssen, W. E.; Jaspars, M.; Naismith, J. H. Structural Analysis of Leader Peptide Binding Enables Leader-Free Cyanobactin Processing. Nat Chem Biol 2015, 11, 558–563. https://doi.org/10.1038/nchembio.1841.

(10) Grove, T. L.; Himes, P. M.; Hwang, S.; Yumerefendi, H.; Bonanno, J. B.; Kuhlman, B.; Almo, S. C.; Bowers, A. A. Structural Insights into Thioether Bond Formation in the Biosynthesis of Sactipeptides. J. Am. Chem. Soc. 2017, 139 (34), 11734–11744. https://doi.org/10.1021/jacs.7b01283.

(11) Latham, J. A.; Iavarone, A. T.; Barr, I.; Juthani, P. V.; Klinman, J. P. PqqD Is a Novel Peptide Chaperone That Forms a Ternary Complex with the Radical S-Adenosylmethionine Protein PqqE in the Pyrroloquinoline Quinone Biosynthetic Pathway. J. Biol. Chem. 2015, 290, 12908–12918. https://doi.org/10.1074/jbc.M115.646521.

(12) Mavaro, A.; Abts, A.; Bakkes, P. J.; Moll, G. N.; Driessen, A. J. M.; Smits, S. H. J.; Schmitt, L. Substrate Recognition and Specificity of the NisB Protein, the Lantibiotic Dehydratase Involved in Nisin Biosynthesis. J. Biol. Chem. 2011, 286 (35), 30552–30560. https://doi.org/10.1074/jbc.M111.263210.

(13) Mitchell, D. A.; Lee, S. W.; Pence, M. A.; Markley, A. L.; Limm, J. D.; Nizet, V.; Dixon, J. E. Structural and Functional Dissection of the Heterocyclic Peptide Cytotoxin Streptolysin S. J. Biol. Chem. 2009, 284 (19), 13004–13012. https://doi.org/10.1074/jbc.M900802200.

(14) Regni, C. A.; Roush, R. F.; Miller, D. J.; Nourse, A.; Walsh, C. T.; Schulman, B. A. How the MccB Bacterial Ancestor of Ubiquitin E1 Initiates Biosynthesis of the Microcin C7 Antibiotic. EMBO J 2009, 28 (13), 1953–1964. https://doi.org/10.1038/emboj.2009.146.

(15) van der Donk, W. A.; Nair, S. K. Structure and Mechanism of Lanthipeptide Biosynthetic Enzymes. Current opinion in structural biology 2014, 29, 58–66. https://doi.org/10.1016/j.sbi.2014.09.006.

(16) Johnson, M.; Zaretskaya, I.; Raytselis, Y.; Merezhuk, Y.; McGinnis, S.; Madden, T. L. NCBI BLAST: A Better Web Interface. Nucleic Acids Research 2008, 36 (Web Server), W5–W9. https://doi.org/10.1093/nar/gkn201.

(17) Finn, R. D.; Bateman, A.; Clements, J.; Coggill, P.; Eberhardt, R. Y.; Eddy, S. R.; Heger, A.; Hetherington, K.; Holm, L.; Mistry, J.; Sonnhammer, E. L.; Tate, J.; Punta, M. Pfam: The Protein Families Database. Nucleic acids research 2014, 42, D222–D230. https://doi.org/10.1093/nar/gkt1223.

(18) Klinman, J. P.; Bonnot, F. Intrigues and Intricacies of the Biosynthetic Pathways for the Enzymatic Quinocofactors: PQQ, TTQ, CTQ, TPQ, and LTQ. Chem. Rev. 2014, 114 (8), 4343–4365. https://doi.org/10.1021/cr400475g.

(19) Evans, R. L.; Latham, J. A.; Xia, Y.; Klinman, J. P.; Wilmot, C. M. Nuclear Magnetic Resonance Structure and Binding Studies of PqqD, a Chaperone Required in the Biosynthesis of the Bacterial Dehydrogenase Cofactor Pyrroloquinoline Quinone. Biochemistry 2017, 56 (21), 2735–2746. https://doi.org/10.1021/acs.biochem.7b00247.

(20) Finn, R. D.; Coggill, P.; Eberhardt, R. Y.; Eddy, S. R.; Mistry, J.; Mitchell, A. L.; Potter, S. C.; Punta, M.; Qureshi, M.; Sangrador-Vegas, A.; Salazar, G. A.; Tate, J.; Bateman, A. The Pfam Protein Families Database: Towards a More Sustainable Future. Nucleic Acids Res. 2016, 44, D279–285. https://doi.org/10.1093/nar/gkv1344.

(21) Soding, J.; Biegert, A.; Lupas, A. N. The HHpred Interactive Server for Protein Homology Detection and Structure Prediction. Nucleic Acids Res 2005, 33 (Web Server), W244–W248. https://doi.org/10.1093/nar/gki408.

(22) McGuffin, L. J.; Bryson, K.; Jones, D. T. The PSIPRED Protein Structure Prediction Server. Bioinformatics 2000, 16 (4), 404–405. https://doi.org/10.1093/bioinformatics/16.4.404.

(23) Tietz, J. I.; Schwalen, C. J.; Patel, P. S.; Maxson, T.; Blair, P. M.; Tai, H.-C.; Zakai, U. I.; Mitchell, D. A. A New Genome-Mining Tool Redefines the Lasso Peptide Biosynthetic Landscape. Nat. Chem. Biol. 2017, 13, 470–478. https://doi.org/10.1038/nchembio.2319.

(24) DiCaprio, A. J.; Firouzbakht, A.; Hudson, G. A.; Mitchell, D. A. Enzymatic Reconstitution and Biosynthetic Investigation of the Lasso Peptide Fusilassin. J. Am. Chem. Soc. 2019, 141 (1), 290–297. https://doi.org/10.1021/jacs.8b09928.

(25) Hudson, G. A.; Burkhart, B. J.; DiCaprio, A. J.; Schwalen, C. J.; Kille, B.; Pogorelov, T. V.; Mitchell, D. A. Bioinformatic Mapping of Radical *S* -Adenosylmethionine-Dependent Ribosomally Synthesized and Post-Translationally Modified Peptides Identifies New Cα, Cβ, and Cγ-Linked Thioether-Containing Peptides. J. Am. Chem. Soc. 2019, jacs.9b01519. https://doi.org/10.1021/jacs.9b01519.

(26) Schwalen, C. J.; Hudson, G. A.; Kille, B.; Mitchell, D. A. Bioinformatic Expansion and Discovery of Thiopeptide Antibiotics. J. Am. Chem. Soc. 2018, 140 (30), 9494–9501. https://doi.org/10.1021/jacs.8b03896.

(27) Gerlt, J. A.; Bouvier, J. T.; Davidson, D. B.; Imker, H. J.; Sadkhin, B.; Slater, D. R.; Whalen, K. L. Enzyme Function Initiative-Enzyme Similarity Tool (EFI-EST): A Web Tool for Generating Protein Sequence Similarity Networks. Biochimica et biophysica acta 2015, 1854, 1019–1037. https://doi.org/10.1016/j.bbapap.2015.04.015.

(28) Su, G.; Morris, J. H.; Demchak, B.; Bader, G. D. Biological Network Exploration with Cytoscape Current protocols in bioinformatics / editoral board, Andreas D. Baxevanis … [et al.] 2014, 47, 8 13 1–24. https://doi.org/10.1002/0471250953.bi0813s47.

(29) Finn, R. D.; Clements, J.; Arndt, W.; Miller, B. L.; Wheeler, T. J.; Schreiber, F.; Bateman, A.; Eddy, S. R. HMMER Web Server: 2015 Update. Nucleic acids research 2015, 43, W30–W38. https://doi.org/10.1093/nar/gkv397.

(30) Altschul, S. Gapped BLAST and PSI-BLAST: A New Generation of Protein Database Search Programs. Nucleic Acids Res. 1997, 25 (17), 3389–3402. https://doi.org/10.1093/nar/25.17.3389.

(31) UniProt, C. UniProt: A Hub for Protein Information. Nucleic acids research 2015, 43, D204–D212. https://doi.org/10.1093/nar/gku989.

(32) Zhang, Q.; Doroghazi, J. R.; Zhao, X.; Walker, M. C.; van der Donk, W. A. Expanded Natural Product Diversity Revealed by Analysis of Lanthipeptide-like Gene Clusters in Actinobacteria. Applied and environmental microbiology 2015, 81, 4339–4350. https://doi.org/10.1128/aem.00635-15.

(33) Walker, M. C.; Eslami, S. M.; Hetrick, K. J.; Ackenhusen, S. E.; Mitchell, D. A.; van der Donk, W. A. Precursor Peptide-Targeted Mining of More than One Hundred Thousand Genomes Expands the Lanthipeptide Natural Product Family. 2019, submitted for publication.

(34) Mirdita, M.; von den Driesch, L.; Galiez, C.; Martin, M. J.; Söding, J.; Steinegger, M. Uniclust Databases of Clustered and Deeply Annotated Protein Sequences and Alignments. Nucleic Acids Res. 2017, 45 (D1), D170–D176. https://doi.org/10.1093/nar/gkw1081.

(35) Blin, K.; Medema, M. H.; Kottmann, R.; Lee, S. Y.; Weber, T. The AntiSMASH Database, a Comprehensive Database of Microbial Secondary Metabolite Biosynthetic Gene Clusters. Nucleic acids research 2016. https://doi.org/10.1093/nar/gkw960.

(36) Steinegger, M.; Söding, J. MMseqs2 Enables Sensitive Protein Sequence Searching for the Analysis of Massive Data Sets. Nat. Biotechnol. 2017, 35 (11), 1026–1028. https://doi.org/10.1038/nbt.3988.

(37) Remmert, M.; Biegert, A.; Hauser, A.; Söding, J. HHblits: Lightning-Fast Iterative Protein Sequence Searching by HMM-HMM Alignment. Nat. Methods 2011, 9 (2), 173–175. https://doi.org/10.1038/nmeth.1818.

(38) Medema, M. H.; Kottmann, R.; Yilmaz, P.; Cummings, M.; Biggins, J.; de Bruijn, I.; Chooi, Y. H.; Claesen, J.; Coates, R. C.; Cruz-Morales, P.; Duddela, S.; Duesterhus, S.; Edwards, D.; Fewer, D. P.; Garg, N.; Geiger, C.; Gomez-Escribano, J. P.; Greule, A.; Hadjithomas, M.; Haines, A. S.; Helfrich, E. J.; Ishida, K.; Jones, A. C.; Jones, C. S.; Jungmann, K.; Kegler, C.; Kim, H. U.; Koetter, P.; Krug, D.; Masschelein, J.; Melnik, A. V.; Mantovani, S. M.; Monroe, E.; Moore, M.; Moss, N.; Nützmann, H. W.; Pan, G.; Pati, A.; Petras, D.; Reen, J.; Rosconi, F.; Rui, Z.; Tian, Z.; Tobias, N. J.; Tsunematsu, Y.; Wiemann, P.; Wickoff, E.; Yan, X.; Yim, G.; Yu, F.; Xie, Y.; Aigle, B.; Apel, A. K.; Balibar, C. J.; Balskus, E.; Barona-Gomez, F.; Bechthold, A.; Bode, H. B.; Borriss, R.; Brady, S.; Brakhage, A.; Caffrey, P.; Cheng, Y.-Q.; Clardy, J.; Cox, R.; De Mot, R.; Donadio, S.; Donia, M. S.; van der Donk, W. A.; Dorrestein, P. C.; Doyle, S.; Driessen, A.; Ehling-Schulz, M.; Entian, K. D.; Fischbach, M. A.; Gerwick, L.; Gerwick, W. H.; Gross, H.; Gust, B.; Hertweck, C.; Höfte, M.; Jensen, S. E.; Ju, J.; Katz, L.; Kaysser, L.; Klassen, J.; Keller, N. P.; Kormanec, J.; Kuipers, O. P.; Kuzuyama, T.; Kyrpides, N.; Kwon, H. J.; Lautru, S.; Lavigne, R.; Lee, C.; Linquan, B.; Liu, X.; Liu, W.; Luzhetskyy, A.; Mahmud, T.; Mast, Y.; Méndez, C.; Metsä-Ketelä, M.; Mitchell, D.; Moore, B. S.; Moreira, L. M.; Müller, R.; Neilan, B.; Nett, M.; Nielsen, J.; O’Gara, F.; Oikawa, H.; Osbourn, A.; Osburne, M.; Ostash, B.; Payne, S.; Pernodet, J. L.; Petricek, M.; Piel, J.; Ploux, O.; Raaijmakers, J. M.; Salas, J. A.; Schmitt, E. K.; Scott, B.; Seipke, R. F.; Shen, B.; Sherman, D.; Sivonen, K.; Smanski, M.; Sosio, M.; Süssmuth, R.; Tahlan, K.; Thomas, C. M.; Tang, Y.; Truman, A. W.; Viaud, M.; Walton, J.; Walsh, C. T.; Weber, T.; van Wezel, G.; Wilkinson, B.; Willey, J.; Wohlleben, W.; Wright, G.; Ziemert, N.; Zhang, C.; Zotchev, S.; Breitling, R.; Takano, E.; Glöckner, F. O. The Minimum Information about a Biosynthetic Gene Cluster (MIBiG) Specification. Nature chemical biology 2015, In revision.

(39) Bushin, L. B.; Clark, K. A.; Pelczer, I.; Seyedsayamdost, M. R. Charting an Unexplored Streptococcal Biosynthetic Landscape Reveals a Unique Peptide Cyclization Motif. J. Am. Chem. Soc. 2018, 140 (50), 17674–17684. https://doi.org/10.1021/jacs.8b10266.

(40) Hamada, T.; Matsunaga, S.; Yano, G.; Fusetani, N. Polytheonamides A and B, Highly Cytotoxic, Linear Polypeptides with Unprecedented Structural Features, from the Marine Sponge, *Theonella s Winhoei*. J. Am. Chem. Soc. 2005, 127 (1), 110–118. https://doi.org/10.1021/ja045749e.

(41) Zhang, Z.; Hudson, G. A.; Mahanta, N.; Tietz, J. I.; van der Donk, W. A.; Mitchell, D. A. Biosynthetic Timing and Substrate Specificity for the Thiopeptide Thiomuracin. J. Am. Chem. Soc. 2016, 138, 15511–15514. https://doi.org/10.1021/jacs.6b08987.

(42) Cheung, W. L.; Chen, M. Y.; Maksimov, M. O.; Link, A. J. Lasso Peptide Biosynthetic Protein LarB1 Binds Both Leader and Core Peptide Regions of the Precursor Protein LarA. ACS Cent. Sci. 2016, 2, 702–709. https://doi.org/10.1021/acscentsci.6b00184.

(43) Dunbar, K. L.; Tietz, J. I.; Cox, C. L.; Burkhart, B. J.; Mitchell, D. A. Identification of an Auxiliary Leader Peptide-Binding Protein Required for Azoline Formation in Ribosomal Natural Products. J. Am. Chem. Soc. 2015, 137 (24), 7672–7677. https://doi.org/10.1021/jacs.5b04682.

(44) Haft, D. H.; Selengut, J. D.; Richter, R. A.; Harkins, D.; Basu, M. K.; Beck, E. TIGRFAMs and Genome Properties in 2013. Nucleic Acids Res. 2013, 41, D387–D395. https://doi.org/10.1093/nar/gks1234.

(45) Khusainov, R.; Moll, G. N.; Kuipers, O. P. Identification of Distinct Nisin Leader Peptide Regions That Determine Interactions with the Modification Enzymes NisB and NisC. FEBS open bio 2013, 3, 237–242. https://doi.org/10.1016/j.fob.2013.05.001.

(46) Schramma, K. R.; Bushin, L. B.; Seyedsayamdost, M. R. Structure and Biosynthesis of a Macrocyclic Peptide Containing an Unprecedented Lysine-to-Tryptophan Crosslink. Nat. Chem. 2015, 7, 431–437. https://doi.org/10.1038/nchem.2237.

(47) Morinaka, B. I.; Lakis, E.; Verest, M.; Helf, M. J.; Scalvenzi, T.; Vagstad, A. L.; Sims, J.; Sunagawa, S.; Gugger, M.; Piel, J. Natural Noncanonical Protein Splicing Yields Products with Diverse β-Amino Acid Residues. Science 2018, 359 (6377), 779–782. https://doi.org/10.1126/science.aao0157.

(48) Zhang, Z.; van der Donk, W. A. Nonribosomal Peptide Extension by a Peptide Amino-Acyl TRNA Ligase. J. Am. Chem. Soc. 2019, 141 (50), 19625–19633. https://doi.org/10.1021/jacs.9b07111.

