## Supplemental Information for "RRE-Finder: A Genome-Mining Tool for Class-Independent RiPP Discovery"

Address: 600 South Mathews Avenue, Urbana, Illinois 61801, USA.

Address: Droevendaalsesteeg 1, 6708 PB Wageningen, The Netherlands

Address: Sylviusweg 72, 2333 BE, Leiden, The Netherlands.

ORCID IDs:

Douglas A Mitchell: 0000-0002-9564-0953

Marnix H. Medema: 0000-0002-2191-2821

Gilles P. van Wezel: 0000-0003-0341-1561

**Table of Contents:**

Experimental Methods ..... S3
Figure S1: Structural homology and sequence divergence of the RRE ..... S8
Table S1: Prevalence of the RRE domain within RiPP classes ..... S9
Table S2: Representative RRE domains with published structures ..... S10
Figure S2: Sequence similarity network of PF05402 (PqqD) ..... S11
Figure S3: Example RiPP biosynthetic gene clusters for precision mode models ..... S12
Table S3: Description of RRE-containing proteins targeted by precision mode ..... S13
Table S4: RRE-Finder analysis times compared to HHPred ..... S14
Table S5: Model validation of precision mode for select RiPP classes ..... S15
Table S6: Validation of RRE-Finder modes against MIBiG database ..... S16
Table S7: Exploratory mode false positives in non-RiPP biosynthetic gene clusters ..... S17
Figure S4: Structural homology of the RRE to DNA-binding elements ..... S18
Figure S5: RRE-containing proteins found in type II PKS clusters ..... S19
Figure S6: Sequence similarity network of retrieved UniProt proteins by phylogeny ..... S20
Figure S7: Sequence similarity network of retrieved UniProt proteins by bit score ..... S21
Figure S8: Overlap of retrieved UniProt proteins in most populous RiPP classes ..... S22
Figure S9: Overlap of retrieved UniProt proteins in YcaO/RRE-dependent RiPP classes ..... S23
Table S8: Conservation of α3 and β3 regions of the RRE ..... S24
Figure S10: Representative phylogenetic tree for retrieved UniProt proteins ..... S25
Table S9: RRE-containing proteins in UniProt found only by exploratory mode ..... S26
Figure S11: Example RiPP biosynthetic gene clusters found by RRE-Finder ..... S27
Table S10: Description of RRE-containing proteins found by RRE-Finder ..... S28
Supporting References ..... S29

### Experimental Methods

**Generation of precision mode models.** Precision mode was generated to accurately predict the presence of RRE domains specific to characterized RiPP classes, as well as RRE domains in select bioinformatically-predicted RRE-dependent RiPP clusters. There are 28 models employed by precision mode of RRE-Finder, each specific to a given discrete or fused RRE protein within a characterized RiPP class (see Figure S3 for represented classes). Each precision mode model consists of a custom profile hidden Markov model (pHMM). To build each HMM, 5-20 representative sequences were selected from a given RRE class for seed sequence alignment. For several RiPP classes, an extensive bioinformatic survey of biosynthetic gene clusters has been conducted. When available, these datasets were employed to select seed sequences, which included: lanthipeptides[1], lasso peptides[2], thiopeptides[3], cyanobactins[4], bottromycins[5], linear azol(in)e-containing peptides [LAPs, including heterocycloanthracins, plantazolicins, nitrile hydratase-like leader peptides (NHLP)-derived RiPPs, Nif11-derived RiPPs, goadsporins, and cytolysins][6], pantocins/microcins[7], and radical *S*-adenosylmethionine-derived RiPPs (including sactipeptides, ranthipeptides, QhpD, and streptides)[8-9]. In these cases, sequence diversity was evaluated by generating a sequence similarity network (SSN) using the Enzyme Function Initiative Enzyme Similarity Tool (EFI-EST) tool[10] and visualizing the SSN with Cytoscape[11]. Five to twenty sequences (depending on number of clusters in the SSN) were selected from divergent clusters on the SSN.

Bioinformatic datasets were not available for the following RRE-dependent RiPP classes: PQQ[12], proteusins, mycofactocins, trifolitoxins, spliceotides, and 3-thiaglutamate. In these cases, a list of homologous sequences to a canonical gene were obtained with position iterative BLAST (PSI-BLAST)[13] with three iterations and a cutoff of 0.05 in November 2019 using GenBank. Once a list of homologous sequences was obtained, an SSN was generated in the same manner as above and diverse sequences were selected for seed sequence alignment.

Seed sequences were analyzed for the presence of an RRE domain using the HHPred webtool (https://toolkit.tuebingen.mpg.de)[14]. A protein was considered to contain an RRE if part or all of the protein matched a PqqD model (either PDB 5SXY or 3G2B) with 80% probability or greater. All proteins containing RRE domains were excised *in silico* to contain only the residues matching the relevant PqqD model. Excised RRE sequences were then aligned using MAFFT 7.450[15]. MAFFT alignments were run using the L-INS-I alignment option on a computer running macOS Mojave version 10.14.6. Multiple sequence alignments were used directly to generate a pHMM using HMMER version 3.3[16]. Models were built using the hmmbuild function and pressed into binary form using the hmmpress function.

**Validation of precision mode models.** Precision mode models were validated against the full datasets from which seed sequences were chosen, excluding the sequences which were included in the pHMMs themselves. For each model, the pHMM was run against the full dataset for the relevant RiPP class using the hmmscan function of HMMER3.3[16]. Hmmscan was run with a bit score cutoff of 25 with all other options set to default. A given model was deemed functional if >95% of RRE-containing protein sequences in a dataset were retrieved by the pHMM at this bit score threshold. In cases where this criterion was not met, sequences not retrieved by the model were used to enrich the original seed sequence alignment and an improved model was generated. In cases where an extensive bioinformatic survey was not available for a certain RiPP class, model accuracy was assessed in two ways: First, the set of homologous proteins generated by PSI-BLAST during model generation was tested against the pHMM using hmmscan with a bit score cutoff of 25. Second, an hmmsearch was performed using the HMMER3.3 webtool (https://www.ebi.ac.uk/Tools/hmmer/search) against the UniProtKB database. The biosynthetic gene clusters surrounding gene hits were visualized using the RODEO webtool[2] (http://rodeo.scs.illinois.edu). A model was considered valid if >95% of the dataset of homologs were hit by the model *and* >90% of proteins hit in the UniProtKB database co-occurred with genes belonging to Pfams known to associate with that RiPP class. Finally, all models were tested for false-positive rates. All models were run against a dataset of 3,000 protein sequences selected from across the datasets used for generating all precision mode models using hmmscan at a bit score cutoff of 35. Models were considered to have acceptably low false positive rates if <100 hits for any given model belonged to a divergent RiPP class.

As described in the main text, precision mode models were also validated against a set of ~5,000 proteins from the antiSMASH database. These protein sequences were employed in the generation of exploratory mode, and thus were a form of cross-validation between the two modes of RRE-Finder. This dataset consists of RRE-containing proteins primarily from the thiopeptide, lasso peptide, lanthipeptide, sactipeptide, and LAP classes. Not all proteins contained within the dataset canonically contain RRE domains, particularly those belonging to class II-IV lanthipeptides. All precision mode models were assessed by hmmscan searches against this dataset with bit score cutoffs of 15, 25, and 35 (representing low, medium and high stringencies).

**Generation of exploratory mode.** Exploratory mode was generated for the purpose of identifying RRE sequences with higher divergence from RREs in known RiPP classes in a more unbiased manner than precision mode. For exploratory mode, we constructed a truncated version of the HHpred pipeline originally used for the detection of RREs[14]. In this pipeline, a query sequence is first expanded with HHblits into a multiple sequence alignment (MSA) using a database of interest[17]. The secondary structure of the MSA is predicted using the adds.pl script available in the HHsuite tool, for which it uses PSIPRED[18]. The MSA is then searched with HHsearch against a second database, which consisted of three sequences from the Protein Databank (PDB) describing crystal structures of reference RRE domains (PDB IDs: 5V1T, 5SXY, 3G2B). To closely mimic the original method for RRE detection, we used the uniclust30 database for MSA generation, which is also used for the HHPred pipeline (version from August 2018, available from https://uniclust.mmseqs.com). This database contains all sequences from the UniProt database clustered with MMseqs2[19] at a cutoff of 30% pairwise sequence identity for MSA generation.

For the initial generation of an RRE database, we used the above pipeline to search 5,000 RiPP BGCs from the antiSMASH database with the uniclust30 database. Regions showing distant similarity to the reference RRE domains (probability ≥40%, length ≥50 residues) were extracted with 15 flanking residues on each side, and the extracted regions were resubmitted to the same pipeline with a higher cutoff to confirm the results (probability ≥90%, length ≥50 residues). Several RREs were added for the LAP and streptide RiPP families, of which no entries were available in the antiSMASH database.

The resulting database of RREs was used for generation of a custom HHpred database as described in the documentation of the HHsuite tool, including the addition of secondary structure predictions with PSIPRED. In parallel, all RREs found were clustered with MMSeqs2 using default settings (pairwise identity ≥80%) and the sequences in each cluster of RREs were aligned using MUSCLE[20]. The resulting alignment was converted into .a3m format using the reformat.pl script available in the HHsuite tool. Each alignment was then further enriched with more homologous sequences from the UniProt database by using HHblits with the uniclust30 database with three iterations. Finally, the expanded alignments were converted into pHMMs using HMMER3.3.

In exploratory mode, each query is first subjected to hmmsearch using the pHMMs described above. Queries passing the cutoff (see main text) and with minimum alignment length of 50 residues have the relevant regions extracted, including 15 flanking residues on each side. The candidate RRE region is then subjected to the same HHpred pipeline described above. In the first step of MSA generation, however, the custom database containing RRE regions is used instead of the uniclust30 database. RRE regions showing significant homology to the reference RRE domains (length ≥50 and probability ≥90%) are considered hits.

**Reducing false positives.** To remove sequences containing transcriptional regulators (a large source of false-positives using the exploratory mode methodology), we constructed a list of Pfam pHMMs containing a variety of DNA-binding regulators and other helix-turn-helix domains that share distant structural homology to the RRE domain. Each resulting hit is searched against this database with hmmsearch using the trusted cutoffs of each pHMM. Overlap of a regulator with a found RRE is indicated in the output file.

**Analysis of MIBiG database.** The pipeline described above was used to analyze all proteins from the MIBiG database (version 1.4), using bit score cutoffs ranging from 15 to 50. The resulting hits were separated into those belonging to RiPP and non-RiPP BGCs. Hits from the RiPP BGCs were additionally clustered per RiPP class. RiPP BGCs containing only precursors were removed.

**Analysis of UniProtKB database.** The pipeline described above was used to analyze all proteins from the UniProtKB/TrEMBL database (UniProt release 2019_09). A bit score cutoff of 25 was used for precision mode and the initial filter of exploratory mode. For exploratory mode, proteins identified as likely regulators were removed before analyzing the queries with the RRE-Finder pipeline.

For the discovery of new classes, UniProt hits found by both modes of RRE-Finder, in particular using the auxiliary models of precision mode, were annotated with Pfam (version 32.0)[21]. Several hits containing a Pfam domain that indicated an enzymatic activity were selected, and their surroundings were investigated, as well as their overlap with antiSMASH. In addition, the presence of RRE domains in these hits was confirmed by submitting them through the HHPred pipeline (https://toolkit.tuebingen.mpg.de/tools/hhpred).

For analysis of the UniProtKB database using precision mode, the HMMER3.3 webtools were used. Each model was individually run through hmmsearch of the UniProtKB database with a bit score cutoff of 25. Hit results for each model were compiled and duplicate protein accessions were removed to determine the exact number of unique protein hits for precision mode of RRE-Finder. Information on duplicate hits from two or more precision mode models were used to determine model overlap and RRE relatedness, as shown in Figures S10 and S11.

**Generation of sequence similarity networks and diversity-maximized phylogenetic tree.** The unique protein accessions from hmmsearch of the UniProtKB database using precision mode were directly used to generate a SSN using the EFI-EST[10] webtool (https://efi.igb.illinois.edu/efi-est/) and visualized with Cytoscape[11]. All sequences were excised to consist of only the RRE domain using a custom script. This script employs hmmsearch to identify the residues of a protein hit corresponding the query pHMM and includes only those residues in the FASTA output. All SSNs shown are either a RepNode60 or RepNode80 network, meaning that protein sequences sharing greater than 60 or 80% sequence identity are conflated into one node on the network. In general, alignment scores for network visualization were chosen to reflect a cutoff where sequences with >40% sequence identity cluster together. For the networks shown in this work, these alignment scores were 18 and 22 (representative of E-value cutoffs of $10^{-18}$ and $10^{-22}$, respectively).

A diversity-maximized, maximum likelihood phylogenetic tree was generated by first selecting a smaller subset of the sequences represented on the SSN. All sequences represented by clusters consisting of 1-3 nodes were included in the tree. For larger clusters, a random sampling of 10% of the sequences in the cluster was used for tree generation. All sequences were excised to contain only the RRE using the same methods as above. The subset of sequences was used to generate a multiple sequence alignment using MAFFT 7.450[15]. MAFFT alignments were run using the L-INS-I alignment option. The MSA was transformed into an approximate-maximum-likelihood tree using FastTree 2.1[22] with the default Jones-Taylor-Thornton (JTT) model. The tree was visualized using the Interactive Tree of Life (iTOL) website (http://itol.embl.de/).

**Integration of RRE-Finder into RODEO and antiSMASH.** Precision mode models have also been incorporated into both the GitHub and webtool versions of RODEO 2. Included is an option to score RRE domains, which if selected, will show which precision mode models are matched along with the default Pfam matches. The integration of precision mode is in-progress for version 6.0 of antiSMASH, which is currently in the development phase and will be reported elsewhere. In addition, the standalone tool available on GitHub (https://github.com/antismash) will be capable of detecting RREs in precision mode and exploratory mode directly from antiSMASH output. RRE-Finder is available as a standalone command-line tool (https://github.com/Alexamk/RREFinder).

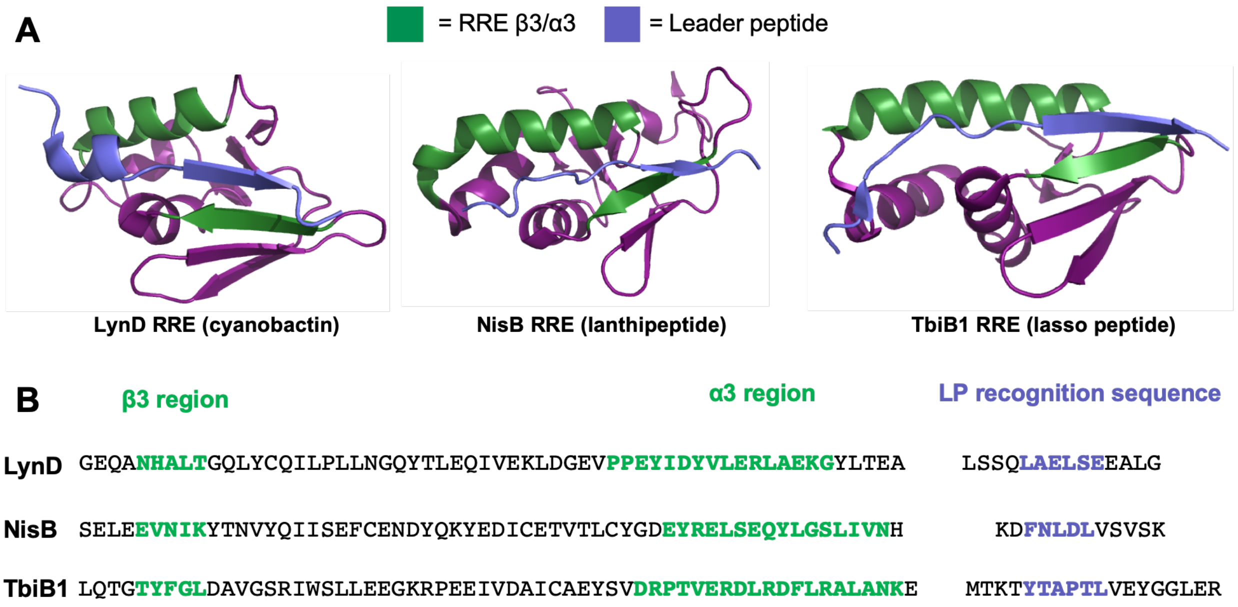


**Figure S1. Structural homology and sequence divergence of the RRE.** (A) The crystal structures of three RRE domains (excised for LynD and NisB) are shown from three RiPP classes. The leader peptide is highlighted in blue, while the conserved cleft in the RRE that binds the leader peptide (LP) is highlighted in green. (B) The truncated protein sequences of each RRE are shown, with the same regions responsible for substrate binding highlighted.

**Table S1. Prevalence of the RRE domain within RiPP classes.** RRE domains are present in over 50% of RiPP classes produced by prokaryotes. These classes are listed in the table along with information pertaining to the type of RRE fusion present and the class-defining modification(s). The example product listed is an archetypal example of the class, and not necessarily the first discovered member of this class. The year of BGC discovery refers to the year in which a product of this class was determined to be a RiPP. In cases where a BGC for more than one member of a class was discovered (e.g. thiopeptides), a representative literature citation is given. Note that ranthipeptides were reclassified as such in 2017, but these RiPPs had been previously bioinformatically characterized and classified as SCIFFs. LAP = linear azol(in)e-containing peptides.

| Class Name | Example Product | Class-Defining Modification and/or Enzyme | RRE Type | Phylogenetic Distribution | Year of BGC Discovery | Citation DOI |
|---|---|---|---|---|---|---|
| Lanthipeptides | Nisin A | Lanthionines (Cys linkage to Ser/Thr β-carbon) | Fused to LanB protein (class I lanthipeptides only) | Archaea and bacteria | 1988 | doi.org/10.1038/333276a0 |
| Pyrroloquinoline quinones (PQQ) | PQQ cofactor | Oxidative cyclization of Glu and Tyr, rSAM installed | Discrete RRE | Bacteria | 1989 | 10.1128/jb.171.1.447-455.1989 |
| Lasso peptides | Microcin J25 | Macrolactam ring with C-terminus threaded through | Fused to leader peptidase or discrete | Archaea and bacteria | 1996 | 10.1128/jb.178.12.3661-3663.1996 |
| LAPs | Microcin B17 | Thiazol(in)es and (methyl)oxazol(in)es, YcaO-dependent | Fused to YcaO domain or stand-alone/E1-like | Archaea and bacteria | 1996 | 10.1126/science.274.5290.1188 |
| Sactipeptides | Subtilosin | Sactionines (Cα linked thioether), rSAM installed | Fused to rSAM sactionine enzyme | Bacteria, primarily firmicutes | 2000 | 10.1128/JB.182.11.3266-3273.2000 |
| Pantocins/ Microcins | Pantocin A | Claisen condensation of glutamic acid residues | Fused | Bacteria | 2003 | doi.org/10.1002/anie.200351054 |
| Cyanobactins | Patellamide A | Protease with PatA homology | Fused to YcaO domain (azoline-containing cyanobactins only) | Cyanobacteria | 2008 | doi.org/10.1038/nchembio.84 |
| Thiopeptides | Thiostrepton | [4+2] cycloaddition | Fused to the F-component of the cyclodehydratase | Bacteria, primarily actinobacteria and firmicutes | 2009 | 10.1073/pnas.0900008106 |
| Mycofactocins | Mycofactocin | Crosslinking of Val and Tyr, rSAM installed | Discrete RRE | Actinobacteria (especially *Mycobacterium*) | 2011 | doi.org/10.1186/1471-2164-12-21 |
| Bottromycins | Bottromycin A1 | Macrolactamidine, YcaO-dependent | Fused to rSAM methyltransferase | Actinobacteria | 2012 | doi.org/10.1039/C2SC21190D |
| Proteusins | Polytheonamide | Nitrile hydratase-derived leader peptide | Fused to rSAM epimerase and rSAM methyltransferase | Bacteria | 2012 | 10.1126/science.1226121 |
| Streptides | Streptide | Trp-Lys crosslinked, rSAM installed | Fused to rSAM enzyme | Mostly firmicutes (especially *Streptococcus*) | 2015 | 10.1038/nchem.2237 |
| Ranthipeptides | Freyrasin | Radical, non-Cα linked thioether, rSAM installed | Fused to rSAM enzyme | Bacteria, primarily firmicutes | 2017 | 10.1021/jacs.9b01519 |
| α-Keto β-amino acid-containing peptides | PlpA | Alpha keto amide linkages, rSAM installed | Discrete RRE | Archaea and bacteria | 2018 | 10.1126/science.aao0157 |
| Rotapeptides | TQQ | Radical oxygen-to-alpha-carbon-linked peptides | Fused to rSAM enzyme | Bacteria, primarily firmicutes | 2019 | 10.1021/jacs.9b05151 |
| Ryptides | RRR | Arg-Tyr crosslinked, rSAM installed | Fused to rSAM enzyme | Bacteria | 2019 | 10.1021/jacs.9b09210 |

**Table S2. Representative RRE domains with published structures.** Representative RRE domains that have been structurally characterized are listed below. LAP = linear azol(in)e-containing peptides. PQQ = pyrroloquinoline quinone.

| Protein | RiPP Class | PDB Accession | UniProtKB Accession | Citation DOI |
|---|---|---|---|---|
| LynD | Cyanobactin | 4V1T | A0YXD2 | 10.1038/nchembio.1841 |
| TruD | Cyanobactin | 4BS9 | B2KYG8 | 10.1002/anie.201306302 |
| NisB | Lanthipeptide | 5WD9 | P20103 | 10.1038/nature13888 |
| McbB | LAP | 6GOS | P23184 | 10.1016/j.molcel.2018.11.032 |
| TfuB1 | Lasso peptide | 6JX3 | Q47AT5 | 10.1021/acschembio.9b00348 |
| TbiB1 | Lasso peptide | 5V1V | D1CIZ5 | 10.1073/pnas.1908364116 |
| MccB | Microcin | 6OM4 | Q47506 | 10.1039/c8sc03173h |
| PaaA | Pantocin | 5FF5 | Q9ZAR3 | 10.1021/jacs.5b13529 |
| PqqD | PQQ | 3G2B/5SXY | Q8P6M8 | 10.1002/prot.22461, 10.1021/acs.biochem.7b00247 |
| CteB | Ranthipeptide | 5WGG | A3DDW1 | 10.1021/jacs.7b01283 |
| SkfB | Sactipeptide | 6EFN | O31423 | 10.1074/jbc.RA118.005369 |
| SuiB | Streptide | 5V1T | A0A0Z8EWX1 | 10.1073/pnas.1703663114 |
| TbtB | Thiopeptide | 6EC7 | D6Y502 | 10.1073/pnas.1905240116 |

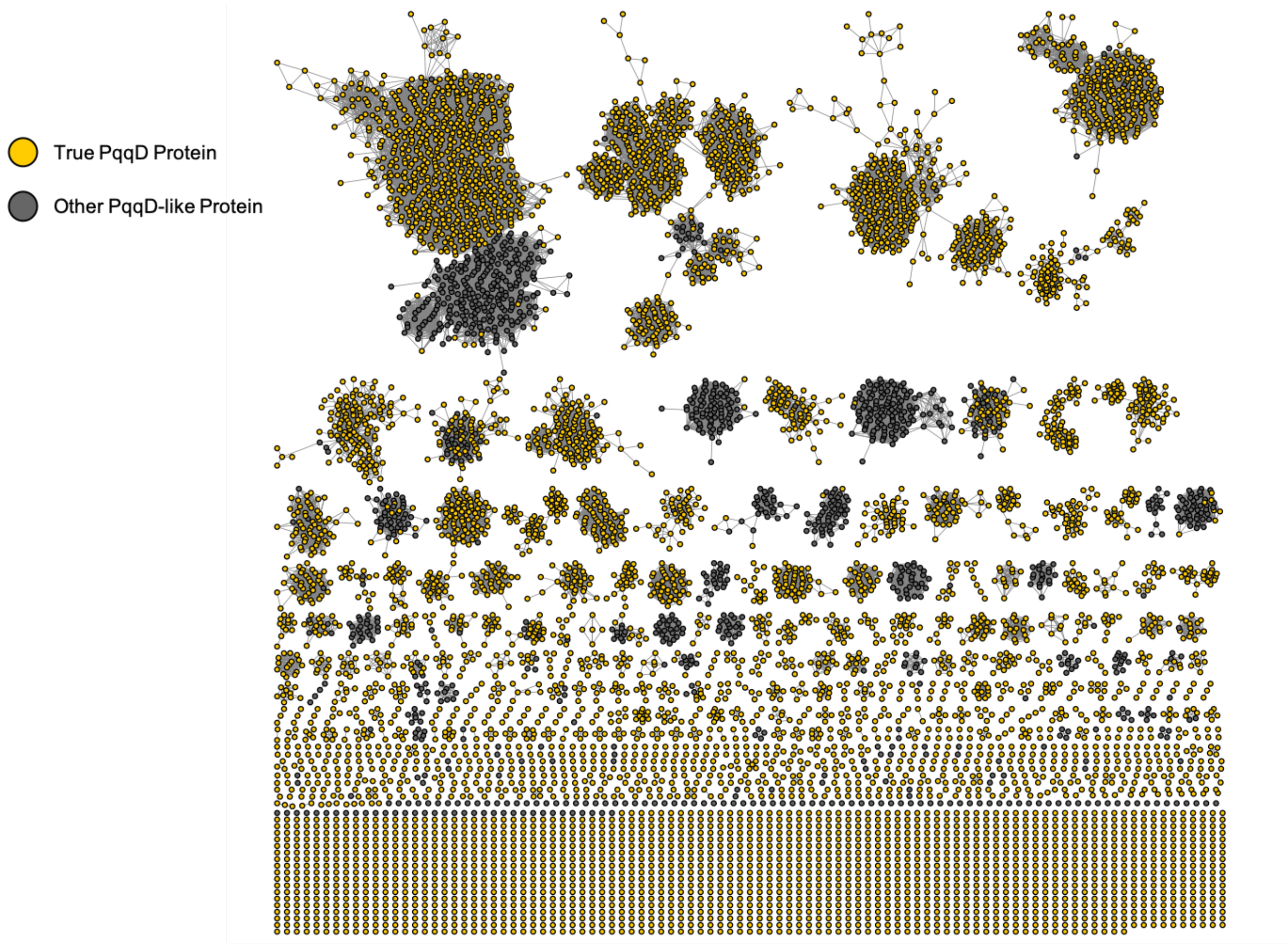


**Figure S2. Sequence similarity network of the PqqD Pfam.** Sequences belonging to PF05402 (PqqD) are represented in the form of a SSN. The network was generated at an alignment score of 25 (E-value = $10^{-25}$) and is presented as a RepNode 80 (protein sequences with greater than 80% identity are conflated to a single node). Nodes are colored in gold if the given protein co-occurs within two genes of a radical SAM enzyme (i.e. a PqqE homolog), indicating that the protein is likely a true PQQ biosynthesis protein. The vast majority of RRE-containing proteins are not retrieved by PF05402.

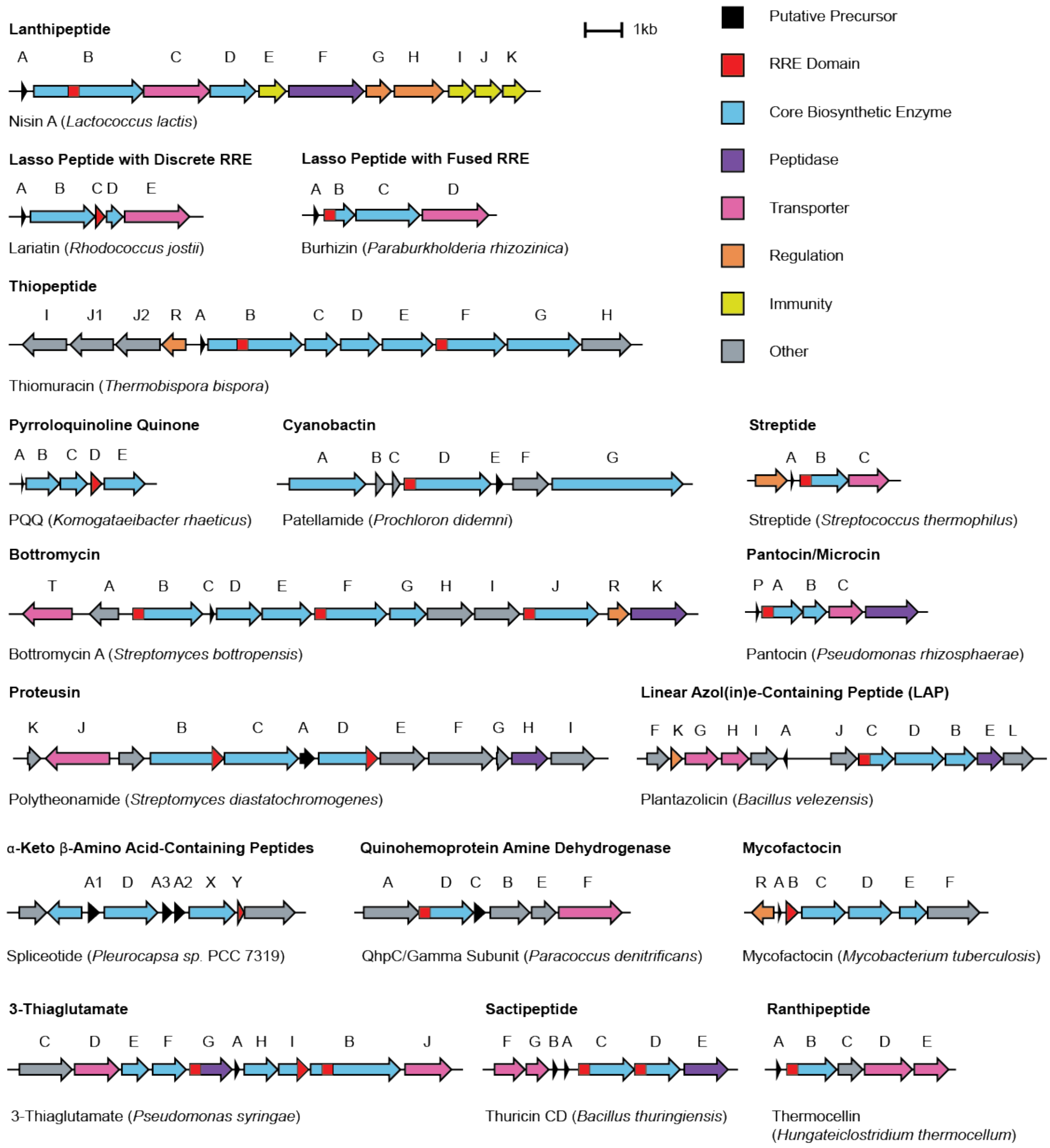


**Figure S3. Representative RiPP gene clusters for precision mode models.** One representative example is given for each RiPP class represented by one or more precision mode models. The relevant class is shown in bold above the BGC, while the specific product encoded by the cluster is shown below the cluster. RRE domains are highlighted in red. In cases where RRE domains are fused to other domains, the red portion of the ORF represents the location of the RRE within the protein. QHNDH = quinohemoprotein amine dehydrogenase. DUF = domain of unknown function. rSAM = radical *S*-adenosyl methionine.

**Table S3. Description of RRE-containing proteins targeted by precision mode.** Further detail is provided for the RRE-containing proteins highlighted in Figure S3. In cases where one BGC contains more than one protein with an RRE, separate NCBI accessions are given for each protein coding sequence.

| Natural Product | Protein | RRE Type | RRE Protein NCBI Accession |
|---|---|---|---|
| Nisin A | NisB | Fused to lanthipeptide dehydratase | ADJ56353.1 |
| Lariatin | LarC | Discrete RRE protein | BAL72548.1 |
| Burhizin | BurB | Fused to lasso peptidase | CBW74825.1 |
| Thiomuracin | TbtB | Fused to lanthipeptide dehydratase | ADG87277.1 |
| Thiomuracin | TbtF | Fused to ocin-ThiF protein | ADG87281.1 |
| PQQ | PqqD | Discrete RRE protein | WP_034930240.1 |
| Patellamide | PatD | Fused to cyclodehydratase | AAY21153.1 |
| Bottromycin | BmbB | Fused to methyltransferase | CCM09442.1 |
| Bottromycin | BmbF | Fused to methyltransferase | CCM09446.1 |
| Bottromycin | BmbJ | Fused to methyltransferase | CCM09450.1 |
| Polytheonamide | PoyB | Fused to methyltransferase | AFS60637.1 |
| Polytheonamide | PoyD | Fused to epimerase | AFS60640.1 |
| Plantazolicin | PznC | Fused to cyclodehydratase | CBJ61638.1 |
| Thuricin CD | TrnC | Fused to rSAM enzyme | AED99784.1 |
| Thuricin CD | TrnD | Fused to rSAM enzyme | AED99785.1 |
| Streptide | SuiB | Fused to rSAM enzyme | ABJ66529.1 |
| Spliceotide | PlpY | Discrete RRE protein | WP_019503879.1 |
| Pantocin | PaaA | Fused to ThiF protein | WP_043190265.1 |
| Thermocellin | CteB | Fused to rSAM enzyme | WP_003517268.1 |
| Mycofactocin | MftB | Discrete RRE protein | WP_019735253.1 |
| QHNDH | QhpD | Fused to rSAM enzyme | SDJ52620.1 |
| 3-Thiaglutamate | PmaB | Fused to short LanB enzyme | KPW26932.1 |
| 3-Thiaglutamate | PmaG | Fused to protease | KPW26903.1 |
| 3-Thiaglutamate | PmaI | Fused to DUF | KPW26921.1 |

**Table S4. RRE-Finder analysis times compared to HHPred.** Both precison and exploratory modes of RRE-Finder significantly decrease analysis times as compared to HHPred, the gold standard for detecting RRE domains. Exploratory mode has longer run times than precision mode, due to the detection of distant protein homology. However, exploratory mode is roughly 250 times faster than HHPred analysis.

| Method | Dataset | Entries | Time Required (h) |
|---|---|---|---|
| RRE-Finder (precision) | MIBiG (all) | 31,025 | 0.002 |
| RRE-Finder (exploratory) | MIBiG (all) | 31,025 | 2.5 |
| HHPred | MIBiG (RiPP only) | 2,513 | 54 |

**Table S5. Model validation of precision mode for select RiPP classes.** Four populous classes of RiPPs were selected for thorough model validation, using the most recent published datasets of predicted BGCs for sactipeptides, ranthipeptides, lanthipeptides, and thiopeptides[1,3,8]. These classes were chosen because of the high quality of the published dataset used for comparison. In all cases, the proteins from each dataset known to contain RRE domains were queried against the relevant precision mode model using HMM scan at low, moderate, and high bit score cutoffs. To determine the false positive rate of the lanthipeptide model, all LanB-type enzymes in the dataset belonging to type II-IV lanthipeptides were queried. These types of lanthipeptides are not predicted to contain RREs, and thus serve as a reasonable false positive. Because the number of class II-IV lanthipeptides is roughly twice that of class I lanthipeptides, the dataset of false positives was much larger for testing this model than for the other three models shown. To determine the false positive rates of the sactipeptide, thiopeptide, and ranthipeptide models, a neighboring protein to each RRE domain was queried. The neighboring proteins queried were ABC transporters (for sactipeptides/ranthipeptides) and cyclodehydratase enzymes (for thiopeptides).

| Dataset | Bit Score | | | Total in Dataset |
|---|---|---|---|---|
| | 15 | 25 | 35 | |
| Lanthipeptide (True Positive) | 1950 | 1910 | 1640 | 2020 |
| Lanthipeptide (False Positive) | 3 | 20 | 90 | 4453 |
| Sactipeptide (True Positive) | 799 | 769 | 690 | 865 |
| Sactipeptide (False Positive) | 1 | 1 | 0 | 865 |
| Ranthipeptide (True Positive) | 2241 | 2150 | 1960 | 2301 |
| Ranthipeptide (False Positive) | 10 | 7 | 4 | 2301 |
| Thiopeptide F Protein (True Positive) | 495 | 492 | 440 | 515 |
| Thiopeptide F Protein (False Positive) | 5 | 3 | 2 | 515 |

**Table S6. Validation of RRE-Finder modes against MIBiG database.** Each detected RRE was grouped by the protein containing it, with an example shown. Precision and exploratory mode combined detect almost all of the hits detected by HHPred (rightmost column). Precision mode readily detects RRE domains in known RiPP classes. Exploratory method detects the same RREs, but additionally detects RREs in recently discovered RiPP BGCs such as the plp and the 3-thiaglutamate gene clusters, as well as in the thioviridamide-like gene clusters and an additional hit (PoyB) in the proteusin gene cluster. Many of these hits were also found by HHPred, highlighting the use for discovery of novel RREs. However, exploratory mode only sparingly detects RREs in the LAP and streptide gene clusters.

| RiPP Class | # curated BGCs | Protein annotation | Example MIBiG Accession | Example Protein | Total | Precision | Exploratory | HHPred |
|---|---|---|---|---|---|---|---|---|
| lasso peptide | 35 | Leader peptidase | BGC0000581 | McjB | 12 | 8 | 10 | 7 |
| | 35 | PqqD-like | BGC0000575 | LarC | 23 | 23 | 23 | 23 |
| lanthipeptide | 31 | LanC-like | BGC001392 | NisC | 1 | 0 | 1 | 1 |
| | 31 | LanB dehydratase | BGC0000535 | NisB | 30 | 29 | 30 | 27 |
| thiopeptide | 24 | Dehydratase | BGC0000613 | TpdB | 21 | 0 | 20 | 6 |
| | 24 | Cyclodehydratase | BGC0000613 | TpdF | 2 | 2 | 2 | 2 |
| | 24 | Radical SAM | BGC0001753 | TbtI | 1 | 0 | 1 | 1 |
| | 24 | ocin_ThiF-like | BGC0000603 | CltD | 23 | 18 | 20 | 17 |
| | 24 | Dehydrogenase | BGC0000613 | TpdE | 5 | 0 | 5 | 3 |
| cyanobactin | 13 | Cyclodehydratase | BGC0000475 | PatD | 8 | 8 | 8 | 8 |
| | 13 | Dehydrogenase | BGC0000475 | PatG | 8 | 0 | 8 | 8 |
| LAP | 10 | Cyclodehydratase | BGC0000569 | PtnD | 7 | 7 | 1 | 3 |
| | 10 | Dehydrogenase | BGC0000565 | GodE | 3 | 1 | 1 | 3 |
| thioamide | 4 | Methyltransferase | BGC0000625 | TvaG | 4 | 0 | 4 | 1 |
| sactipeptide | 4 | Radical SAM | BGC0000600 | ThnB | 5 | 4 | 4 | 4 |
| bottromycin | 4 | Radical SAM | BGC0000468 | BmbB | 12 | 12 | 12 | 0 |
| pheganomycin | 1 | Methyltransferase | BGC0001148 | Pgm4 | 1 | 0 | 1 | 0 |
| | 1 | Radical SAM | BGC0001148 | Pgm3 | 1 | 0 | 1 | 1 |
| proteusin | 1 | Radical SAM | BGC0000598 | PoyB | 1 | 0 | 1 | 1 |
| | 1 | Radical SAM | BGC0000598 | PoyC | 1 | 1 | 1 | 1 |
| | 1 | | BGC0000598 | PoyD | 1 | 1 | 1 | 1 |
| plp | 1 | Radical SAM | BGC0001745 | PlpY | 1 | 1 | 0 | 1 |
| streptide | 1 | Radical SAM | BGC0001209 | SuiB | 1 | 1 | 0 | 0 |
| microcin | 1 | ThiF-like | BGC0000585 | MccB | 1 | 1 | 1 | 1 |
| 3-thiaglutamate | 1 | LanB dehydratase | BGC0001486 | PmaJ | 1 | 1 | 1 | 0 |
| | 1 | | BGC0001486 | PmaI | 1 | 1 | 1 | 1 |
| | 1 | Peptidase | BGC0001486 | PmaG | 1 | 1 | 1 | 1 |

**Table S7. Exploratory mode false positives in non-RiPP BGCs.** Exploratory mode retrieved a total of 51 proteins in non-RiPP BGCs at a bit score cutoff of 25. Many retrieved proteins were transcriptional regulators or proteins with a helix-turn-helix (HTH) motif, which is unsurprising considering structural homology to the RRE domain. Other false positives included several methyltransferase enzymes and proteins with sequence homology to RRE-containing proteins in RiPP clusters. The latter group may contain true positives, as some large non-RiPP BGCs may have poorly defined borders, and therefore contain parts of RiPP gene clusters (e.g. BGC0000696, the gene cluster for gentamicin, contains a LanB dehydratase and a LanC cyclase at the edge of the gene cluster).

| False Positive Type | Number of Proteins Retrieved |
|---|---|
| Transcription Regulators/HTH Domains | 19 |
| Methyltransferases | 12 |
| Associated with Known RiPPs | 15 |
| Other | 12 |

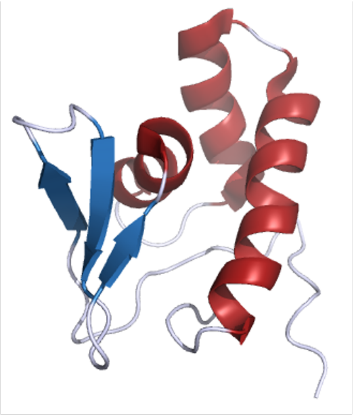

**PqqD NMR Structure**
***Methylobacterium extorquens***
**PDB 5SXY**

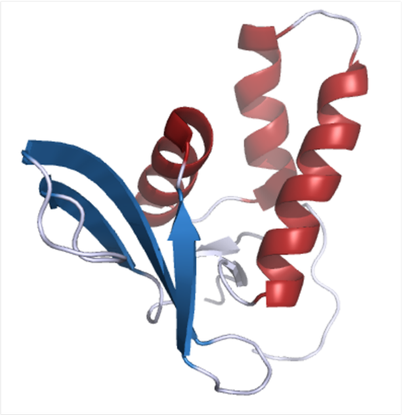

**Transcriptional Regulator**
***Neisseria gonnorrhoeae***
**PDB 3DEE**

**PDB 5SXY**

MEPTAFSGSDVPRLPRGVRLRFDEVRNKHVLLAPERTFDLDDNAVAVLKLVDGRNTVSQIAQILGQTYDADPAIIEADILPMLAGLAQKRVLER

**PDB 3DEE**

HYSNDSKYTPSPAAFIRQYRYDVTHDLQEAETALLIWRNAEDDVMYQTLDGFDMMLLEIMGSSALSFDTLAQTLVEFMPKADNWKNILLGKWSGWIEQRIIIPS

**Figure S4. Structural homology of the RRE to DNA-binding elements.** The RRE consists of a conserved secondary structure of three α-helices and three β-strands, highlighted in blue and red in the structures shown. This secondary structure is also present in many regulatory and DNA-binding elements, such as the truncated DNA-binding portion of the *Neisseria* protein shown. HHPred analysis also shows high structural homology (>90% probability) between several DNA-binding elements and RRE-containing proteins. Sequence similarity between transcription regulators and RRE domains still remains low, with the two sequences shown sharing only 33% sequence identity. Thus, it is plausible that RRE domains evolved from transcriptional regulator proteins.

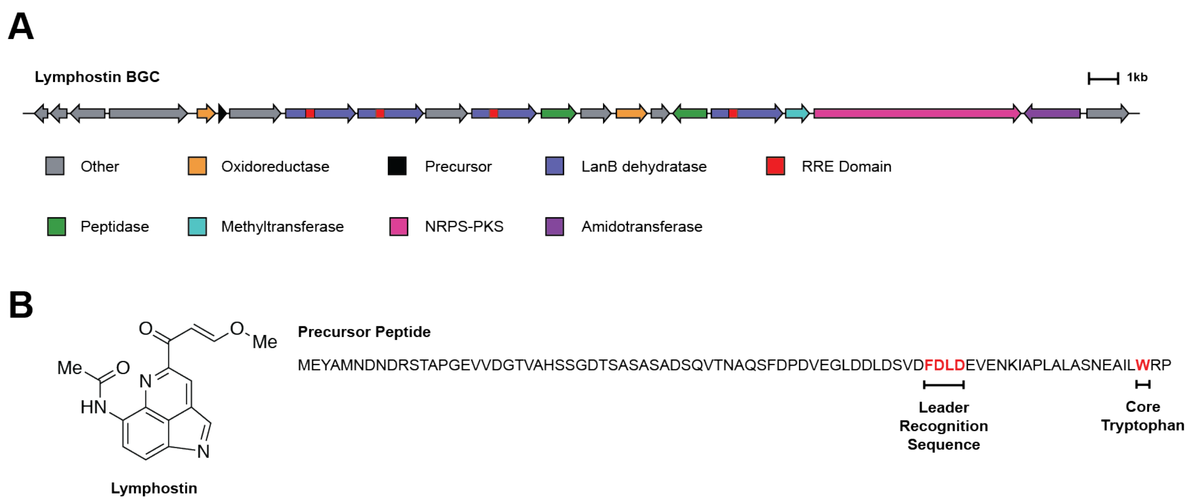


**Figure S5. RRE-containing proteins found in type II PKS clusters.** (A) The biosynthetic gene cluster for lymphostin, a member of the pyrroloquinoline alkaloid class of RiPPs[23]. Many pyrroloquinoline alkaloid (PQA) clusters contain both an NRPS-PKS module as well as one or more LanB-type enzymes containing internal RRE domains. (B) The structure of lymphostin, a RiPP natural product derived from tryptophan.

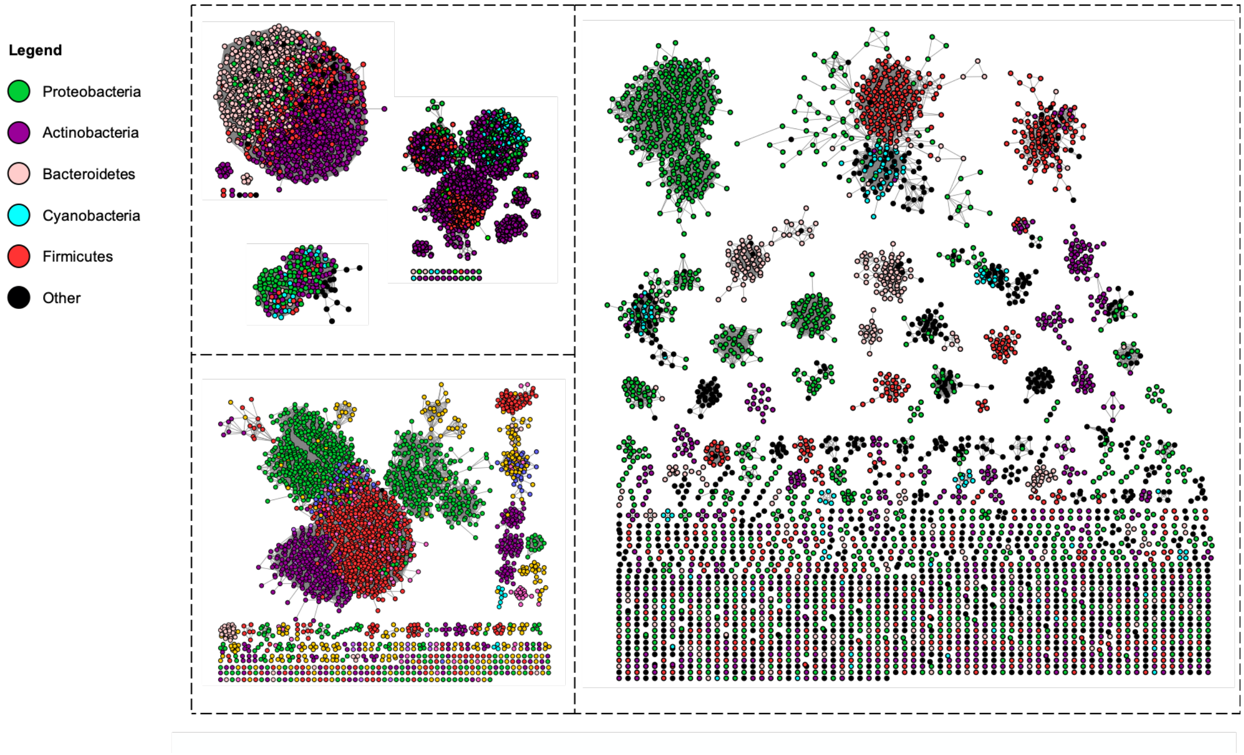


**Figure S6. Sequence similarity network of retrieved UniProt proteins by phylogeny.** The UniProtKB database was searched using all precision mode models at a bit score cutoff of 25. All proteins retrieved are visualized on the sequence similarity network. The SSN is identical to Figure 5 but has been recolored by phylum (alignment score of 22, RepNode60). The SSN was generated using the EFI-EST tool[10] and visualized with Cytoscape[11].

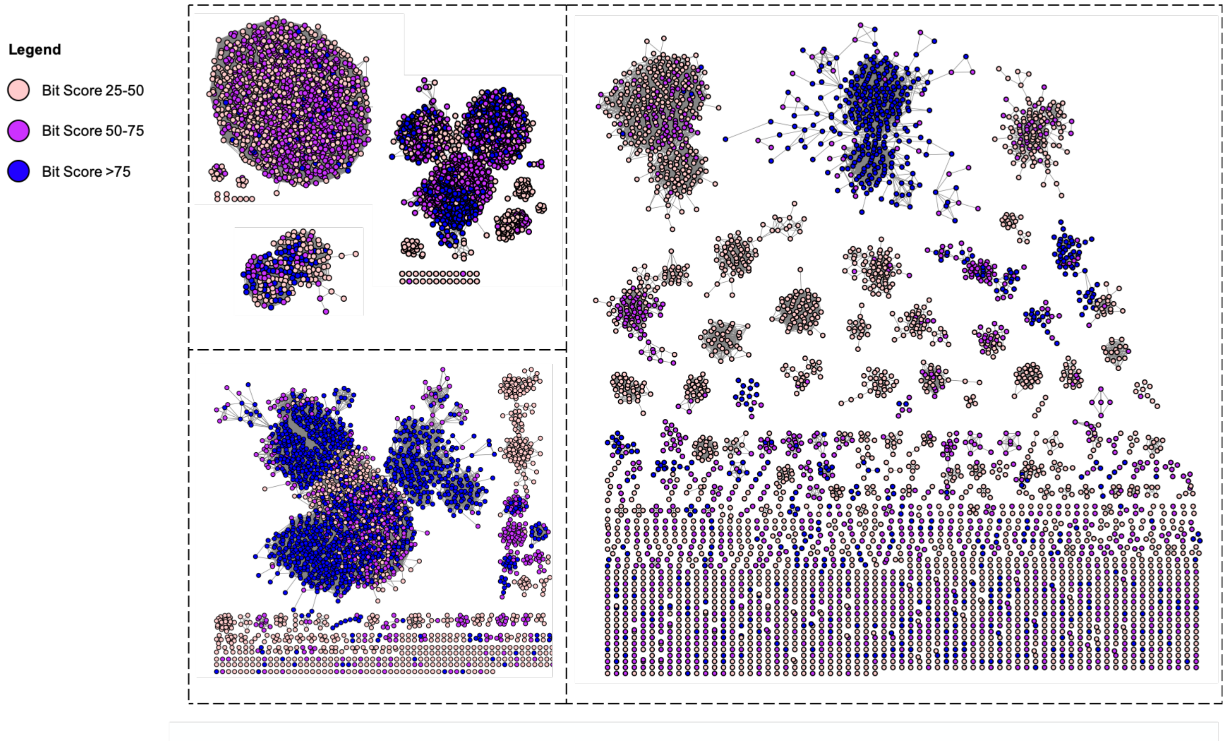


**Figure S7. Sequence similarity network of retrieved UniProt proteins by bit score.** The UniProtKB database was searched using all precision mode models at a bit score cutoff of 25. All proteins retrieved are visualized on the sequence similarity network. All nodes and edges are identical to Figure 5 but are recolored according to the bit score significance of the hit to a precision mode model (alignment score of 22, RepNode60). The network was generated using the EFI-EST tool[10] and visualized with Cytoscape[11].

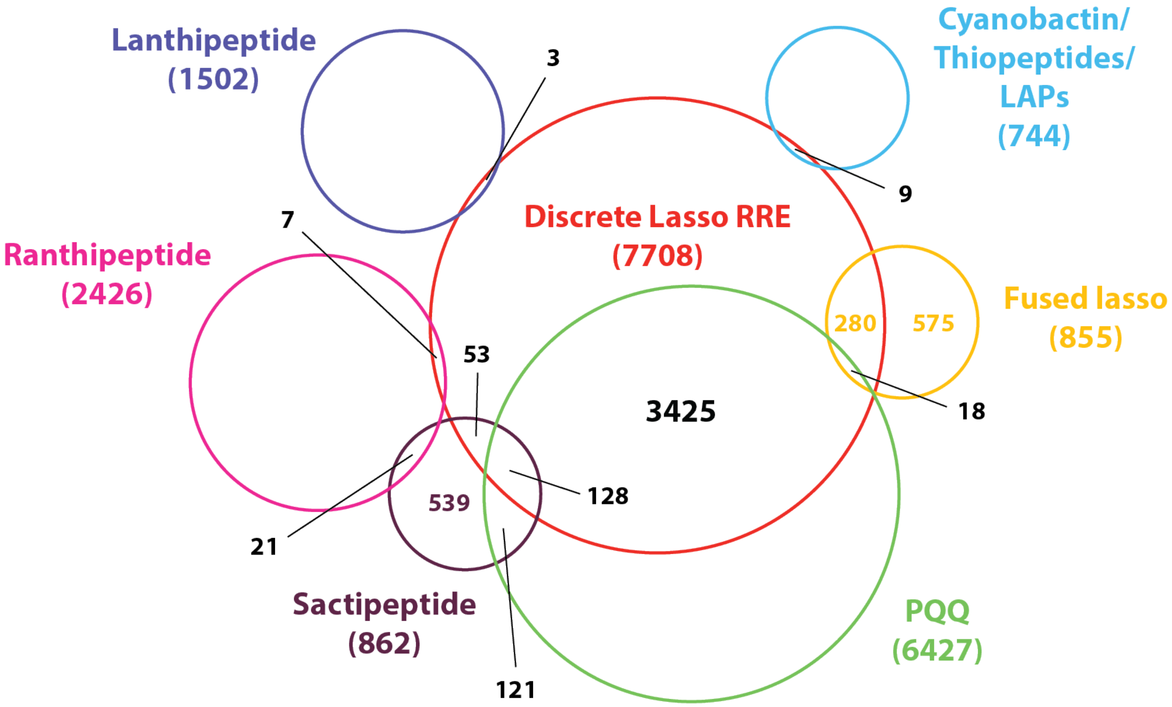


**Figure S8. Overlap of retrieved UniProt proteins in most populous RiPP classes.** Individual precision mode models for the most populous RiPP classes were employed for HMM search of the UniProtKB database at a bit score cutoff of 25. The total number of retrieve sequences for each model is indicated in parentheses. The numbers within circles indicates model redundancy or overlap, owing to the same protein sequence being retrieved by more than one precision mode model. The discrete lasso peptide RRE model retrieves more proteins than anticipated because of similar motifs present in non-lasso clusters. For example, many lasso peptide RREs co-occur in clusters with rSAM enzymes. In addition, there is significant overlap between the RREs of lasso peptides and those from PQQ clusters. Model overlap is similar to what is shown above at a more stringent bit score cutoff of 35.

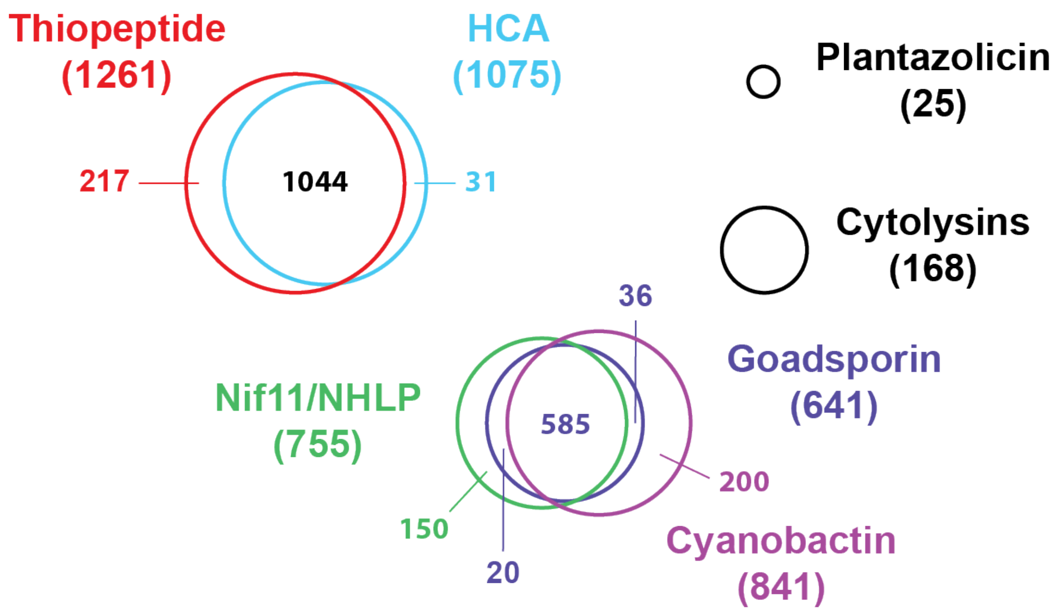


**Figure S9. Overlap of retrieved UniProt proteins in YcaO/RRE-dependent RiPP classes.** Individual precision mode models for RiPP classes containing azole and azoline heterocycles were employed for HMM search of the UniProtKB database at a bit score cutoff of 25. Total number of hits for each model is indicated in parentheses, while numbers within circles indicates model redundancy or overlap, where the same protein sequence was hit by one or more precision mode models. Model overlap reveals that some numbers of retrieved proteins for precision mode are artificially high. For example, there are only ~500 proteins retrieved by the thiopeptide model that co-occur with canonical thiopeptide modifying enzymes, such as the [4+2] cycloaddition enzyme. The other proteins retrieved by this model are heterocycloanthracins, which employ a highly similar leader peptide recognition sequence and RRE domain primary sequence. NHLP, nitrile hydratase-like leader peptide[24]. HCA, heterocycloanthracin.

**Table S8. Conservation of α3 and β3 regions of the RRE.** Residue-level conservation was assessed using three metrics on eight precision mode models. The secondary structures principally responsible for binding the leader peptide (the α3 and β3 regions) were assessed separately from the remainder of the RRE domain. The region of the RRE with the greatest conservation per metric is indicated by red text. Individual RiPP classes were scored by selecting 10 divergent RREs from that class and excising the relevant substructure sequence. For RiPP classes that have significant evolutionary relatedness, a total of 20 sequences were used for the calculations (10 from each class). These data reveal a trend of higher conservation in the α3 and β3 regions of the RRE compared to other regions. Perhaps unsurprisingly, α3 displays the greatest conservation across RiPP classes, given that the contact with the leader peptide is primarily through side chain interactions as opposed to the β3 strand (primarily backbone interactions). HCA, heterocycloanthracin.

| | **Shannon Information Entropy** | | | **ConSurf (0-9 scale)** | | | **AACon (0-9 scale)** | | |
|---|---|---|---|---|---|---|---|---|---|
| | α3 helix | β3 strand | other | α3 helix | β3 strand | other | α3 helix | β3 strand | other |
| Goadsporin | **0.81** | 0.65 | 0.45 | **7** | 6 | 4 | **7** | 6 | 4 |
| Cyanobactin | **0.75** | 0.59 | 0.39 | **7** | 6 | 3 | **7** | 6 | 4 |
| Goadsporin and Cyanobactin | **0.62** | 0.54 | 0.21 | **6** | **6** | 2 | **6** | 5 | 2 |
| Discrete Lasso peptide | **0.43** | 0.33 | 0.23 | **4** | 3 | 2 | **4** | 3 | 2 |
| Fused Lasso peptide | **0.51** | 0.32 | 0.31 | **5** | 3 | 3 | **4** | 3 | 2 |
| Discrete and Fused Lasso peptide | **0.27** | 0.22 | 0.13 | **3** | **3** | 1 | **3** | 2 | 1 |
| Thiopeptide | **0.76** | 0.72 | 0.56 | **7** | **7** | 6 | **7** | **7** | 5 |
| HCA | **0.82** | 0.74 | 0.58 | **8** | 7 | 6 | **8** | 7 | 5 |
| Thiopeptide and HCA | **0.71** | 0.64 | 0.49 | **7** | 6 | 5 | **7** | 6 | 5 |
| Ranthipeptide | **0.68** | 0.57 | 0.42 | **7** | 6 | 4 | **7** | 5 | 4 |
| QhpD | **0.71** | 0.59 | 0.47 | **7** | 6 | 5 | **7** | 6 | 5 |
| Ranthipeptide and QhpD | **0.54** | 0.43 | 0.36 | **5** | 4 | 4 | **5** | 4 | 3 |

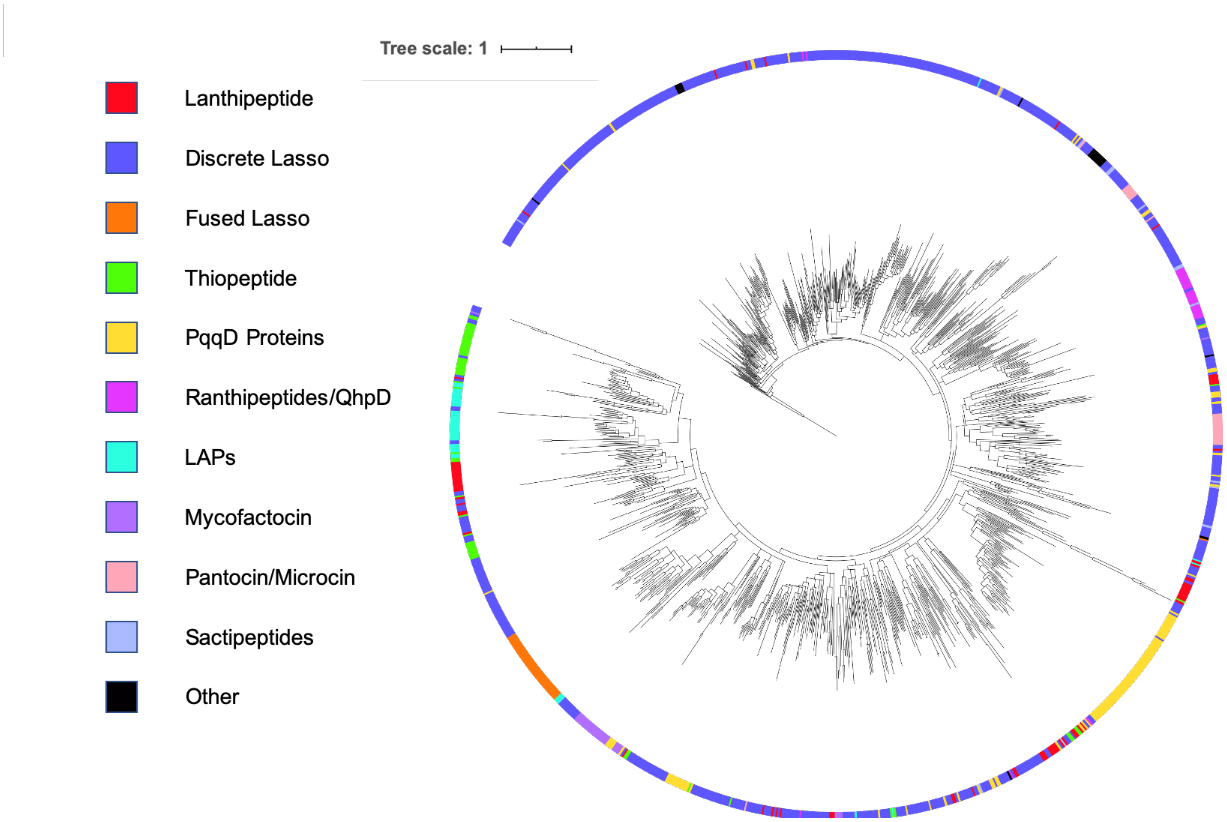


**Figure S10. Representative phylogenetic tree for retrieved UniProt proteins.** Shown are ~20,000 protein sequences retrieved by HMM search of the UniProtKB database using precision mode models. All proteins used for tree generation were truncated *in silico* to contain only the RRE domain. Leaves are colored based on the precision mode model hit with the highest bit score. Given the prevalence of discrete lasso-like RRE domains in non-lasso peptide BGCs, this model shares the most sequence similarity to non-RiPP regulatory proteins, and thus branch most directly from the transcriptional regulator out-group (PDB entry: 3DEE). The tree was generated using FastTree[22] and visualized using the iTol web tool[25].

**Table S9. RRE-containing proteins in UniProt found only by exploratory mode.** Proteins were grouped based on Pfam domains**.** Besides the known RRE-fusions, such as those containing YcaO domains and LanB dehydratase domains, many novel RRE fusions are identified. These include fusions with a wide variety of protein domains not previously linked to RiPP biosynthesis. RRE domain can also be found in a number of small proteins without a protein domain, suggesting these are discrete RREs.

| Protein domain description | Number found |
|---|---|
| DNA-binding proteins or regulators | 23,528 |
| Other (no pfam domain, >= 120 aa) | 14,994 |
| Discrete RRE (no pfam domain, <120 aa) | 2,819 |
| PqqD | 2,410 |
| Metallo beta-lactamase | 1,739 |
| Radical SAMs and iron-sulfur containing proteins | 1,618 |
| LanB | 1,522 |
| Nitroreductase | 1,053 |
| Methyltransferases | 1,050 |
| Memo domain | 475 |
| Oxidoreductases | 123 |
| Translgutaminase | 99 |
| YcaO | 86 |
| Tryptophan halogenase | 81 |
| Tetratricopeptide domain | 77 |
| Cyclic nucleotide binding domain | 73 |
| Peptidase | 68 |
| Cupin domain | 28 |
| Glycosyltransferases | 28 |
| Acetyltransferase | 20 |
| Asparagine synthase | 17 |
| Glutathione S-transferase | 13 |
| LanC | 13 |
| Carbamoyltransferase | 7 |

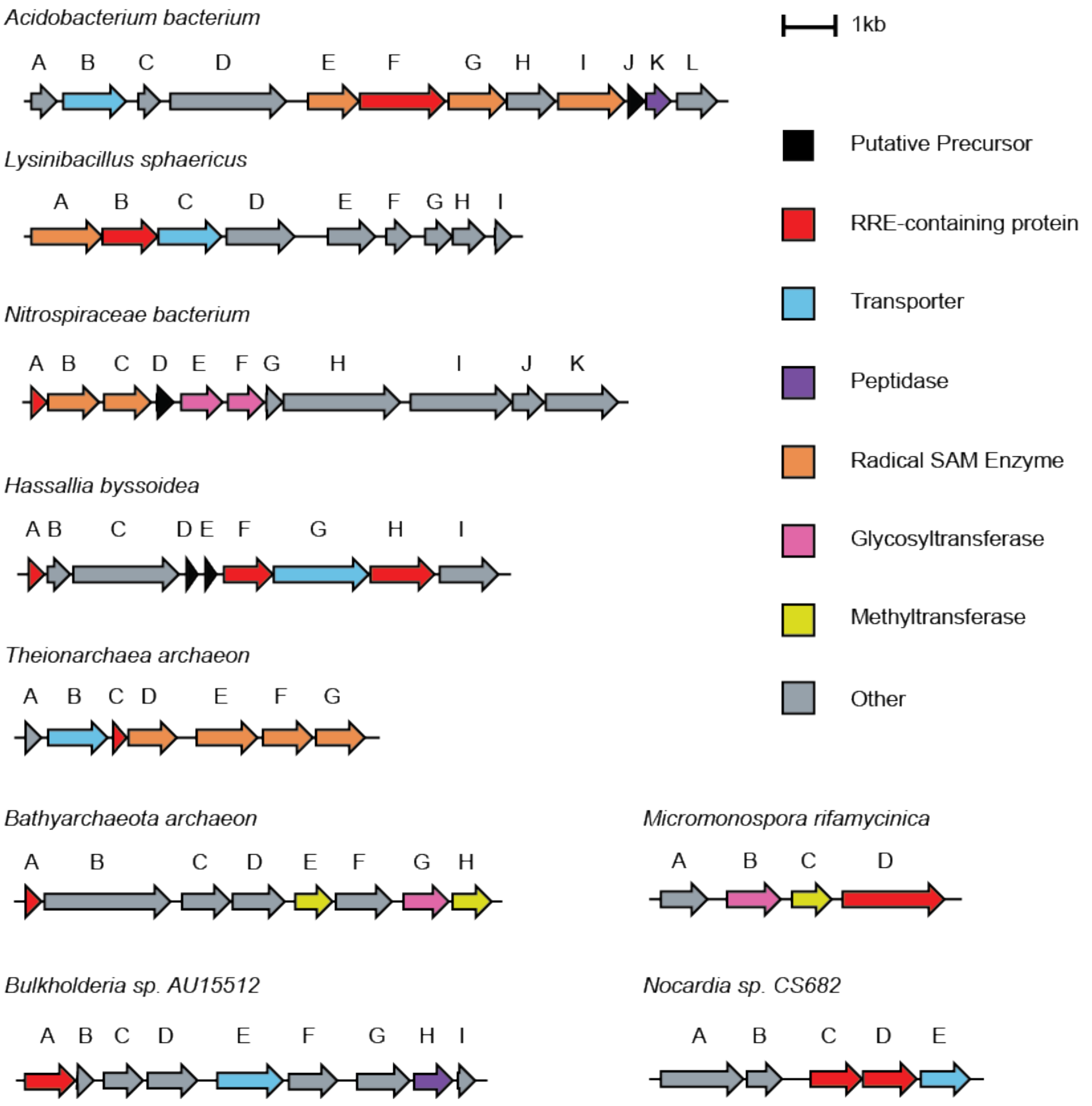


**Figure S11. Example RiPP gene clusters found by RRE-Finder.** Shown are nine BGCs that contain RRE domains in novel contexts. Proteins highlighted in red with asterisks indicate proteins containing RRE domains as predicted by RRE-Finder. All RRE-domain containing proteins are listed in the accompanying table along with protein accessions.

**Table S10. Description of RRE-containing proteins found by RRE-Finder.** Further detail is provided for the RRE-containing proteins shown in Figure S11. The letters used to identify a gene correspond to those used in the biosynthetic gene clusters in Figure S11.

| Organism | Gene | RRE Type | RRE NCBI Accession |
|---|---|---|---|
| *Acidobacterium bacterium* | F | Fused to tetratricopeptide domain | OFW29522.1 |
| *Lysinibacillus sphaericus* | B | Fused to glutathione *S*-transferase | WP_069508305.1 |
| *Nitrospiraceae bacterium* | A | Discrete RRE protein | RPI38387.1 |
| *Hassallia byssoidea* | A | Discrete RRE protein | KIF30015.1 |
| *Hassallia byssoidea* | F | Fused to glycosyltransferase | KIF29242.1 |
| *Hassallia byssoidea* | H | Fused to phosphoribosyl transferase | KIF29244.1 |
| *Theioarchaea archaeon* | C | Discrete RRE protein | KYK35486.1 |
| *Bathyarchaeota archaeon* | A | Discrete RRE protein | OGD46518.1 |
| *Micromonospora rifamycinica* | D | Fused to carbamoyltransferase | WP_067301990.1 |
| *Bulkholderia sp. AU15512* | A | Fused to iron redox enzyme | OXI24931.1 |
| *Nocardia sp. CS682* | C | Fused to heme-oxygenase enzyme | QBS40287.1 |
| *Nocardia sp. CS682* | D | Fused to iron redox enzyme | QBS40286.1 |

### Supplementary References


(1) Walker, M. C.; Eslami, S. M.; Hetrick, K. J.; Ackenhusen, S. E.; Mitchell, D. A.; van der Donk, W. A. Precursor Peptide-Targeted Mining of More than One Hundred Thousand Genomes Expands the Lanthipeptide Natural Product Family. **2019**, submitted for publication.

(2) Tietz, J. I.; Schwalen, C. J.; Patel, P. S.; Maxson, T.; Blair, P. M.; Tai, H.-C.; Zakai, U. I.; Mitchell, D. A. A New Genome-Mining Tool Redefines the Lasso Peptide Biosynthetic Landscape. *Nat. Chem. Biol.* **2017**, *13*, 470–478. https://doi.org/10.1038/nchembio.2319.

(3) Schwalen, C. J.; Hudson, G. A.; Kille, B.; Mitchell, D. A. Bioinformatic Expansion and Discovery of Thiopeptide Antibiotics. *J. Am. Chem. Soc.* **2018**, *140* (30), 9494–9501. https://doi.org/10.1021/jacs.8b03896.

(4) Sardar, D.; Pierce, E.; McIntosh, J. A.; Schmidt, E. W. Recognition Sequences and Substrate Evolution in Cyanobactin Biosynthesis. *ACS Synth. Biol.* **2014**. https://doi.org/10.1021/sb500019b.

(5) Schwalen, C. J.; Hudson, G. A.; Kosol, S.; Mahanta, N.; Challis, G. L.; Mitchell, D. A. In Vitro Biosynthetic Studies of Bottromycin Expand the Enzymatic Capabilities of the YcaO Superfamily. *J. Am. Chem. Soc.* **2017**, *139* (50), 18154–18157. https://doi.org/10.1021/jacs.7b09899.

(6) Cox, C. L.; Doroghazi, J. R.; Mitchell, D. A. The Genomic Landscape of Ribosomal Peptides Containing Thiazole and Oxazole Heterocycles. *BMC Genom.* **2015**, *16*, 778. https://doi.org/10.1186/s12864-015-2008-0.

(7) Ghodge, S. V.; Biernat, K. A.; Bassett, S. J.; Redinbo, M. R.; Bowers, A. A. Post-Translational Claisen Condensation and Decarboxylation En Route to the Bicyclic Core of Pantocin A. *J. Am. Chem. Soc.* **2016**, *138* (17), 5487–5490. https://doi.org/10.1021/jacs.5b13529.

(8) Hudson, G. A.; Burkhart, B. J.; DiCaprio, A. J.; Schwalen, C. J.; Kille, B.; Pogorelov, T. V.; Mitchell, D. A. Bioinformatic Mapping of Radical *S* -Adenosylmethionine-Dependent Ribosomally Synthesized and Post-Translationally Modified Peptides Identifies New Cα, Cβ, and Cγ-Linked Thioether-Containing Peptides. *J. Am. Chem. Soc.* **2019**, jacs.9b01519. https://doi.org/10.1021/jacs.9b01519.

(9) Bushin, L. B.; Clark, K. A.; Pelczer, I.; Seyedsayamdost, M. R. Charting an Unexplored Streptococcal Biosynthetic Landscape Reveals a Unique Peptide Cyclization Motif. *J. Am. Chem. Soc.* **2018**, *140* (50), 17674–17684. https://doi.org/10.1021/jacs.8b10266.

(10) Gerlt, J. A.; Bouvier, J. T.; Davidson, D. B.; Imker, H. J.; Sadkhin, B.; Slater, D. R.; Whalen, K. L. Enzyme Function Initiative-Enzyme Similarity Tool (EFI-EST): A Web Tool for Generating Protein Sequence Similarity Networks. *Biochim. Biophys. Acta* **2015**, *1854*, 1019–1037. https://doi.org/10.1016/j.bbapap.2015.04.015.

(11) Su, G.; Morris, J. H.; Demchak, B.; Bader, G. D. Biological Network Exploration with Cytoscape 3. *Curr Protoc Bioinformatics* **2014**, *47*, 8 13 1-24. https://doi.org/10.1002/0471250953.bi0813s47.

(12) Latham, J. A.; Iavarone, A. T.; Barr, I.; Juthani, P. V.; Klinman, J. P. PqqD Is a Novel Peptide Chaperone That Forms a Ternary Complex with the Radical S-Adenosylmethionine Protein PqqE in the Pyrroloquinoline Quinone Biosynthetic Pathway. *J. Biol. Chem.* **2015**, *290*, 12908–12918. https://doi.org/10.1074/jbc.M115.646521.

(13) Altschul, S. Gapped BLAST and PSI-BLAST: A New Generation of Protein Database Search Programs. *Nucleic Acids Res.* **1997**, *25* (17), 3389–3402. https://doi.org/10.1093/nar/25.17.3389.

(14) Soding, J.; Biegert, A.; Lupas, A. N. The HHpred Interactive Server for Protein Homology Detection and Structure Prediction. *Nucleic Acids Res* **2005**, *33* (Web Server), W244–W248. https://doi.org/10.1093/nar/gki408.

(15) Katoh, K.; Standley, D. M. MAFFT Multiple Sequence Alignment Software Version 7: Improvements in Performance and Usability. *Mol Biol Evol* **2013**, *30*, 772–780. https://doi.org/10.1093/molbev/mst010.

(16) Finn, R. D.; Clements, J.; Arndt, W.; Miller, B. L.; Wheeler, T. J.; Schreiber, F.; Bateman, A.; Eddy, S. R. HMMER Web Server: 2015 Update. *Nucleic acids research* **2015**, *43*, W30–W38. https://doi.org/10.1093/nar/gkv397.
(17) Remmert, M.; Biegert, A.; Hauser, A.; Söding, J. HHblits: Lightning-Fast Iterative Protein Sequence Searching by HMM-HMM Alignment. *Nat Methods* **2012**, *9* (2), 173–175. https://doi.org/10.1038/nmeth.1818.
(18) McGuffin, L. J.; Bryson, K.; Jones, D. T. The PSIPRED Protein Structure Prediction Server. *Bioinformatics* **2000**, *16* (4), 404–405. https://doi.org/10.1093/bioinformatics/16.4.404.
(19) Steinegger, M.; Söding, J. MMseqs2 Enables Sensitive Protein Sequence Searching for the Analysis of Massive Data Sets. *Nat Biotechnol* **2017**, *35* (11), 1026–1028. https://doi.org/10.1038/nbt.3988.
(20) Edgar, R. C. MUSCLE: Multiple Sequence Alignment with High Accuracy and High Throughput. *Nucleic Acids Res* **2004**, *32* (5), 1792–1797. https://doi.org/10.1093/nar/gkh340.
(21) Finn, R. D.; Coggill, P.; Eberhardt, R. Y.; Eddy, S. R.; Mistry, J.; Mitchell, A. L.; Potter, S. C.; Punta, M.; Qureshi, M.; Sangrador-Vegas, A.; Salazar, G. A.; Tate, J.; Bateman, A. The Pfam Protein Families Database: Towards a More Sustainable Future. *Nucleic Acids Res.* **2016**, *44*, D279-285. https://doi.org/10.1093/nar/gkv1344.
(22) Price, M. N.; Dehal, P. S.; Arkin, A. P. FastTree: Computing Large Minimum Evolution Trees with Profiles Instead of a Distance Matrix. *Mol. Biol. Evol.* **2009**, *26* (7), 1641–1650. https://doi.org/10.1093/molbev/msp077.
(23) Miyanaga, A.; Janso, J. E.; McDonald, L.; He, M.; Liu, H.; Barbieri, L.; Eustáquio, A. S.; Fielding, E. N.; Carter, G. T.; Jensen, P. R.; Feng, X.; Leighton, M.; Koehn, F. E.; Moore, B. S. Discovery and Assembly-Line Biosynthesis of the Lymphostin Pyrroloquinoline Alkaloid Family of MTOR Inhibitors in Salinispora Bacteria. *J. Am. Chem. Soc.* **2011**, *133* (34), 13311–13313. https://doi.org/10.1021/ja205655w.
(24) Haft, D. H.; Basu, M. K.; Mitchell, D. A. Expansion of Ribosomally Produced Natural Products: A Nitrile Hydratase- and Nif11-Related Precursor Family. *BMC Biol* **2010**, *8* (1), 70. https://doi.org/10.1186/1741-7007-8-70.
(25) Letunic, I.; Bork, P. Interactive Tree Of Life (ITOL) v4: Recent Updates and New Developments. *Nucleic Acids Res* **2019**, *47* (W1), W256–W259. https://doi.org/10.1093/nar/gkz239.
